# Structural basis of mitochondrial membrane bending by I-II-III2-IV2 supercomplex

**DOI:** 10.1101/2022.06.26.497646

**Authors:** Alexander Mühleip, Rasmus Kock Flygaard, Outi Haapanen, Rozbeh Baradaran, Thomas Gruhl, Victor Tobiasson, Amandine Maréchal, Vivek Sharma, Alexey Amunts

**Affiliations:** Science for Life Laboratory, Department of Biochemistry and Biophysics, Stockholm University, 17165 Solna, Sweden.; Department of Molecular Biology and Genetics, Danish Research Institute of Translational Neuroscience - DANDRITE, Nordic EMBL Partnership for Molecular Medicine, Aarhus University, 8000 Aarhus C, Denmark.; Department of Physics, University of Helsinki, 00014 Helsinki, Finland.; MRC Laboratory of Molecular Biology, Cambridge, United Kingdom.; Institute of Structural and Molecular Biology, Birkbeck College, London, WC1E 7HX, UK.; Institute of Structural and Molecular Biology, University College London, London, WC1E 6BT, UK.; HiLIFE Institute of Biotechnology, University of Helsinki, 00014 Helsinki, Finland.

## Abstract

Mitochondrial energy conversion requires an intricate architecture of the inner mitochondrial membrane^1^. Here we show that in ciliates, the membrane curvature is provided by a supercomplex containing all four respiratory chain components. We report cryo-electron microscopy and cryo-tomography structures of the supercomplex that comprises 150 different proteins and 311 bound lipids, forming a stable 5.8-megadalton assembly. Due to subunit acquisition and extension, complex I associates with a complex IV dimer, generating a wedge-shaped gap that serves as a binding site for complex II. Together with a tilted complex III dimer association, it results in a curved membrane region. Using molecular dynamics simulations, we demonstrate that the divergent supercomplex actively contributes to the membrane curvature induction and cristae tubulation. Our findings explain how the architecture of the native I-II-III_2_-IV_2_ supercomplex reflects the functional specialization of bioenergetics by shaping the membrane.

---

Mitochondrial energy conversion requires an electron transport chain (ETC) that generates a membrane potential across the inner mitochondrial membrane to drive the essential adenosine triphosphate (ATP) formation by F_1_F_o_-ATP synthase. The ETC consists of four multi-subunit membrane complexes: complex I (CI, NADH:ubiquinone oxidoreductase), complex II (CII, succinate:ubiquinone oxidoreductase), complex III (CIII, cytochrome bc1 complex) and complex IV (CIV, cytochrome c oxidase). Structural analyses have shown that these components can organize into supercomplexes containing CI, CIII dimer (CIII_2_), and CIV^1^. CII transfers electrons from succinate via its covalently bound flavin adenine dinucleotide (FAD) and iron-sulfur clusters to ubiquinone (UQ) and is also a component of the TCA cycle, making a functional link between the two central metabolic pathways^2^. Although CII has been suggested to interact with mammalian ETC complexes^3–7^, it was not experimentally found as a part of any characterized supercomplex. In addition, for the bioenergetic process to occur, a specific topology of the cristae membranes that form functionally distinct high-potential compartments is critical^8^. An established mechanism for maintenance of such a topology relies on oligomerization of ATP synthase and its specific interplay with lipids^9–13^. In ciliates, the inner mitochondrial membrane is organized as tubular cristae, which cannot be explained by the helical row assembly of ATP synthase alone^11, 14, 15^.

We purified the intact respiratory supercomplex from the ciliate protist *Tetrahymena thermophila* mitochondria and determined its structure by single-particle cryo–electron microscopy (cryo-EM) (Extended Data Fig. 1 and SI Table 1). At an overall resolution of 2.9 Å, the structure revealed CI, CII, CIII_2_, and CIV dimer (CIV_2_) associated into a 5.8-megadalton supercomplex (Fig. 1). When viewed along the membrane plane, the assembly of more than 300 transmembrane helices displays a bent shape, indicating that the accommodating membrane adopts a local curvature with radius of ∼20 nm (Fig. 1). Focused refinements resolved individual structures that together form an assembly of 150 different protein subunits and 311 bound lipids (Extended Data Figs. 1 and 2, SI Tables 1 and 2). CIV_2_ is associated with the long side of the membrane region of CI, opposite to CIII_2_. This arrangement is markedly different compared to known mammalian supercomplexes^16, 17^ and correlates with the acquisition of four ciliate-specific CI subunits that would clash with the position of CIV as seen in mammals (Fig. 1D, Extended Data Fig. 3, SI Fig. 1). CII is anchored in between CI and CIV, highlighting the unique architecture and composition of the native supercomplex (Fig.1). To further substantiate the occurrence of a functional supercomplex, we performed in-gel activity assays. These confirmed the presence of functional electron-transfer systems in CI, CII and CIV with activities mapping to a common high molecular weight band, which we assign as the intact supercomplex (SI Fig. 2).

**Fig. 1.**
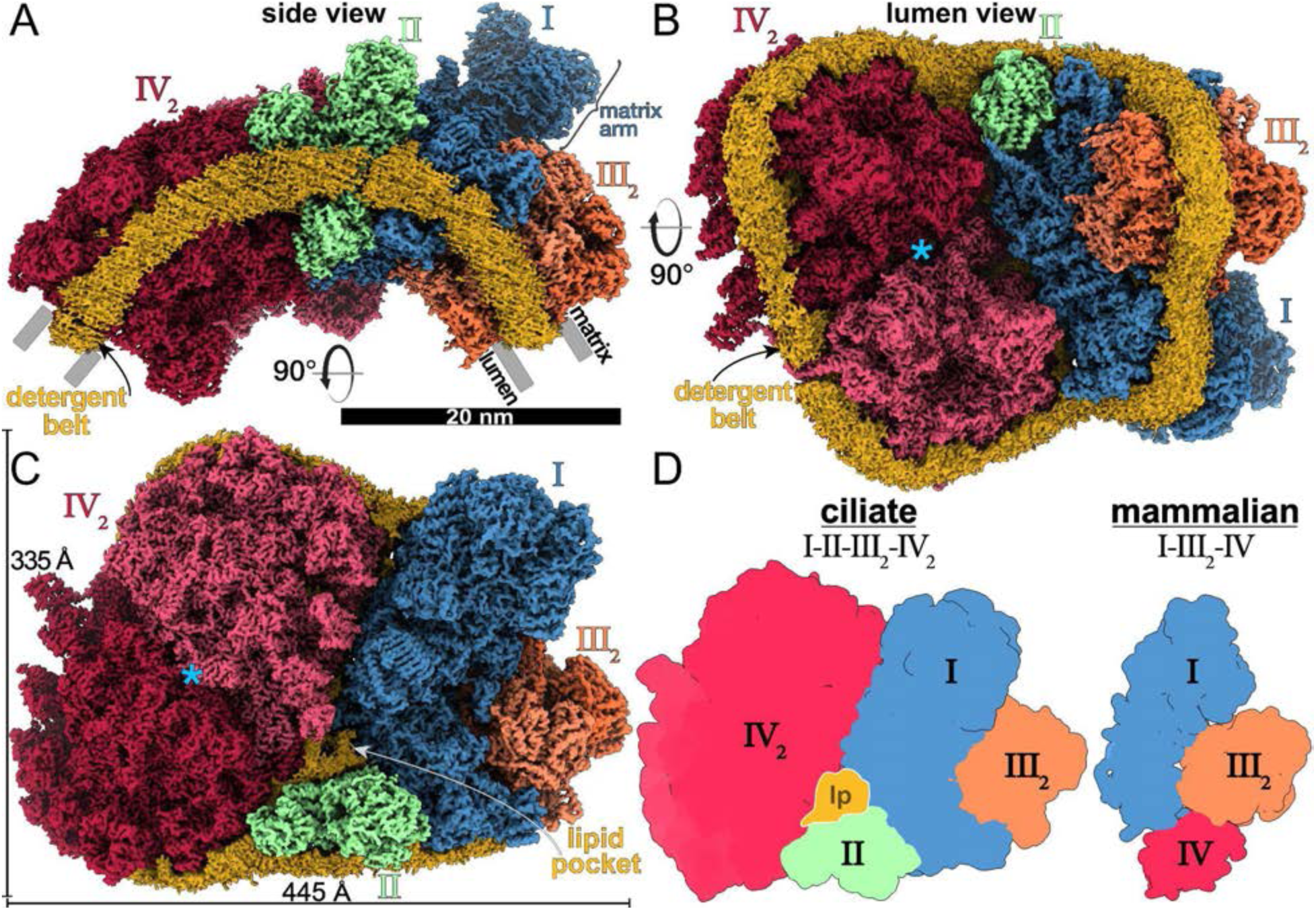
The supercomplex contains all four ETC components. **(A)** Side view of the supercomplex density showing the curved detergent micelle (yellow). **(B)** Lumenal view illustrates how the complexes CI, CII and CIV_2_ stabilize each other. Blue asterisk indicates the symmetry axis of CIV_2_. **(C)** Matrix view shows CII binding in a wedge between CI and CIV_2_, resulting in the enclosure of a lipid pocket (lp). **(D)** Architecture comparison of the ciliate supercomplex (this study) with mammalian respirasome (PDB 5J4Z) highlighting a different location of CIV_2_ that is correlated with acquisition of CI subunits that stabilize CIII_2_.

CIV_2_ is the most divergent of the four ETC complexes (Extended Data Figs. 4 and 5, SI Fig. 3). We modeled 105 lipids, four previously unobserved ubiquinones and 53 protein chains per monomer, of which four are mitochondrially encoded (Extended Data Fig. 2C-F, Extended Data Fig. 6). We found that two of those subunits, previously annotated as ciliate-specific Ymf67 and Ymf68*/*COX3 represent complementary protein fragments, with coding genes split in the mt- genome by tRNA^Trp^ gene insertion (Extended Data Fig. 7). Each fragment has subsequently been extended by over 400 and 200 residues respectively. Together, they form a functional COX3 (COX3a, COX3b), including the conserved seven-TM-helix fold (Extended Data Fig. 7A, SI Fig. 3). In our structure, COX3a and COX3b extend throughout the CIV membrane region, and COX3b has evolved interactions with CI subunits on the matrix side, thereby mediating the supercomplex assembly (Fig. 2, A and B, Extended Data Fig. 7A). Particularly, COX3b forms a contact with a peripheral amphipathic helix of NDUCA1, which is part of a zinc-free γ-carbonic anhydrase heterotrimer (γ-CA) (Fig. 2D). The γ-CA was previously reported in viridiplantae and ciliates (both diaphoretickes)^7, 18^, and our structure demonstrates that it acts as a structural scaffold within the supercomplex architecture. Another CI-IV contact at the same site involves COXTT2 with an N-terminal globlin-like domain that interacts with NDUFA3 and was not resolved in the individual CIV_2_ structure^7^, suggesting that it becomes ordered to mediate supercomplex formation (Fig. 2A and D, Extended Data Fig. 6A). Interestingly, the second major interaction site at the CI-IV interface, which is on the lumenal side of the membrane also involves a fragmented protein subunit, this time from CI (Extended Data Fig. 7C and D).

**Fig. 2.**
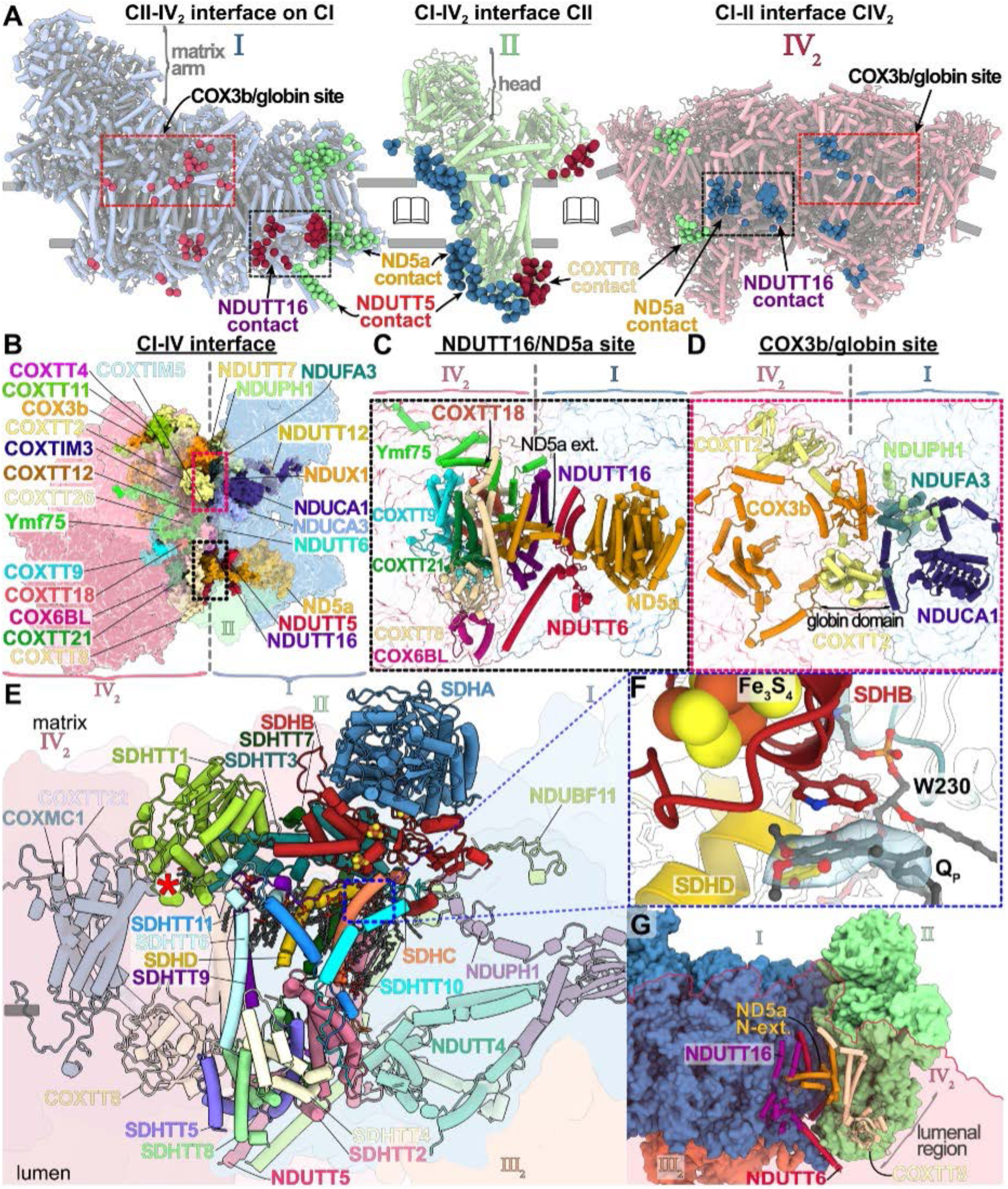
The CI-CIV association, and binding of CII. **(A)** Contact sites of CI with CII-IV_2_ (left), CII with CI-IV_2_ (middle), CIV_2_ with CI-II (right). Interactions are shown as spheres (CI blue, CII green, CIV_2_ red/dark and pink). Only one CIV monomer interacts with CII. Main interaction sites are indicated. **(B)** The CI-IV_2_ interacting subunits are shown in colored surfaces. **(C)** NDUTT16/ND5a contact site. **(D)** COX3b/NDUCA1 contact site. **(E)** CII binding to subunits of CI-IV (transparent), asterisk marks C-type heme. **(F)** CII contains a bound proximal ubiquinone. **(G)** CI, CII and CIV are connected together via the membrane and lumenal regions.

Consistent with the observation with respect to the protein splitting in CIV, here we modeled the N-terminal extension of ND5 fragment (ND5a), as well as the newly identified protein subunit NDUTT16 (from CI) (Fig. 2 A-C). NDUTT16 engages in interactions with at least four subunits of CIV, as well as an interfacial CIV heme group (Fig. 2C).

Our finding of the split core subunits gaining a capacity of establishing inter-complex contacts to stabilise the supercomplex that curves the membrane suggests an evolutionary mechanism by which gene fragmentation, followed by its expansion can convey subunit function. Overall, the CI-IV_2_ interface involves 25 subunits, forming an extensive buried interface of ∼2,300 Å^2^ with a curved membrane region (Fig. 2, A and B).

Tt-CII binds in a wedge-shaped gap formed by CI-CIV in our structure (Fig. 1, A-C). In addition to the four canonical subunits (SDHA-D), it is composed of 11 ciliate-specific subunits SDHTT1-11 (Fig. 2E, Extended Data Fig. 8A). The matrix module SDHA and SDHB forms a conserved head region, containing both the covalently bound FAD and three iron-sulfur clusters (Fig. 2F, Extended Data Fig. 8B). The membrane anchor is formed by two small subunits SDHC (7 kDa) and mitochondria-encoded SDHD (5 kDa), which could only be assigned by locating topologically conserved transmembrane helices in the map (Extended Data Fig. 8 A,C,D). At the lumen, a ∼70-kDa module (SDHTT2, 4, 5, and 8) anchors CII to CI-IV (Fig. 2E, Extended Data Fig. 8A). SDHTT5 interacts with a helix of NDUTT5 protruding from the membrane arm of CI, and with the Surf1-like protein subunit COXTT8 (CIV) (Fig. 2, A and E). Remarkably, at the same position COXTT8 interacts with CI via the N-terminal NDA5a extension, and with Ymf75, COXTT27 and COXTT18 at the CIV dimer interface. Thus, the three complexes are connected together in the lumen (Fig. 2G).

In between SDHB and SDHD, we identified a ligand which we assign as the proximal ubiquinol (Q_p_) (Fig. 2F)^19, 20^. On the matrix side, the 36-kDa soluble subunit SDHTT1 contains a bis-histidine C-type heme group covalently bound by a single cysteine residue (Extended Data Fig. 8E). Although it is exposed to the membrane region, at a distance of ∼60 Å to the Fe_3_S_4_ cluster, the non-canonical heme-c is located too far to participate in direct CII electron transfer (Fig. 2F, Extended Data Fig. 8A,E). To elucidate the presence of additional heme groups in the supercomplex, we recorded absorption spectra of the purified sample (Extended Data Fig. 9). Deconvolution of the merged absorption bands of B- and C-type hemes indicated the presence of at least one additional heme group with absorption at 556 nm.

The presence of a functional ETC with CII is consistent with previous observations that *T. thermophila* can utilize succinate to drive cellular respiration^21^. Our native structure with bound CII, which contributes to the ubiquinol pool, demonstrates that supercomplex assembly is not limited to proton-pumping respiratory chain components (CI, CIII, CIV). Beyond decreasing cytochrome-c transfer distance^22^, this suggests a potential role of supercomplex formation in mediating increased ubiquinone diffusion, as suggested in analogous membrane systems with high protein-lipid ratios^23, 24^. Furthermore, the tubular membrane morphology may require the anchoring of CII into the curved supercomplex to retain it in the functionally relevant cristae, preventing diffusion into flat membrane regions.

CIII_2_ in our structure is tilted with respect to CI by 37° (Fig. 3A and SI Fig. 4). This tilted arrangement offsets the transmembrane region, consistent with its curved membrane environment. The interface involves 20 subunits and 19 bound native lipids interacting through the matrix, transmembrane and luminal sides (Extended Data Fig. 10). When compared to the mammalian counterpart, CIII_2_ is rotated by 41° and shifted ∼14 Å due to acquisition of four CI subunits, as well as the CIII subunit UQCRTT1 (Fig. 3A and Extended Data Fig. 10B,C)^16, 25–27^. This arrangement results in a specific CI-CIII_2_ contact with one copy of Rieske iron sulfur protein (UQCRFS1) interacting with the CI membrane arm (Fig. 3B). The interaction site is further augmented by a hitherto unidentified protein UQCRTT3, which interacts with the luminal head domain of UQCRFS1 and wedges in between the interface to CYC1 (Fig. 3B).

**Fig. 3.**
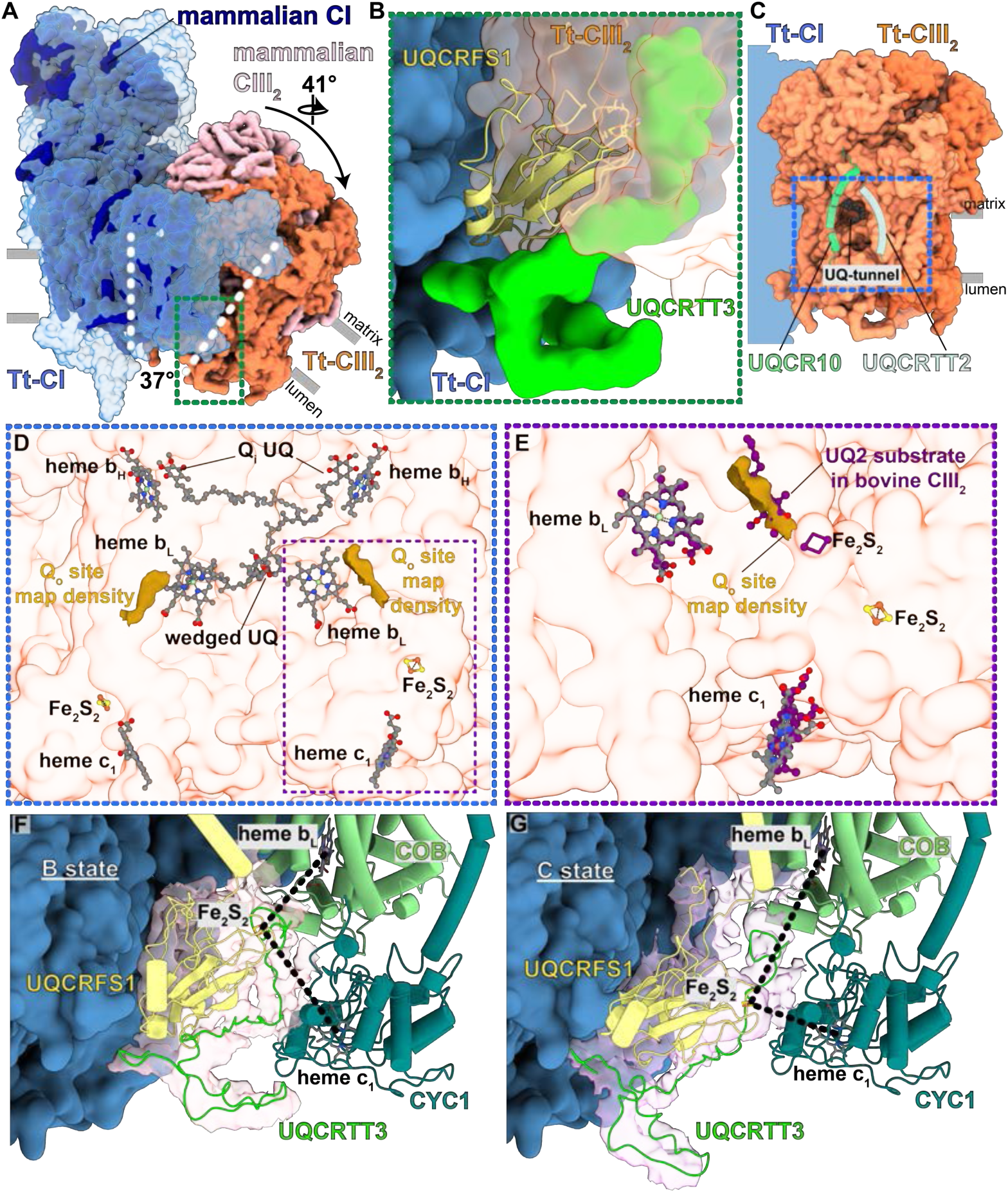
Altered CI-CIII_2_ interface maintains functional symmetry of CIII_2_. **(A)** Superposition of Tt-CI-CIII_2_ with mammalian CI-CIII_2_ (PDB 5J4Z) shows that Tt-CIII_2_ is tilted, rotated and displaced with respect to the CI membrane arm, as indicated by dashed lines and arrows due to acquisition of new proteins. Green box view is shown in B. **(B)** Tilted Tt-CIII_2_ results in an interaction between UQCRFS1 and CI membrane arm together with UQCRTT3. **(C)** Tt-CIII_2_ shows a membrane-accessible tunnel extending to the COB Q_i_ sites. Blue box view is shown in D. **(D)** The Q_i_ ubiquinones are located close to heme b_H_, with one ubiquinone wedged in between the two heme b_L_ molecules. The density map of Tt-CIII_2_ shows two features corresponding to Q_o_ sites. Purple box view is shown in E. **(E)** The Q_o_ site density overlaps with ubiquinol in bovine CIII_2_ structure (PDB 1NTZ). **(F,G)** 3D maps showing B state (F) and C state (G) conformations of UQCRFS1 and UQCRTT3. Only COB and CYC1 proteins are shown for clarity, highlighting distances of Fe_2_S_2_ to heme c_1_ and b_L_ (black dashes).

We traced the membrane-accessible UQ-tunnel, lined by UQCR10 and ciliate-specific subunit UQCRTT2 (Fig. 3C), leading to the COB heme b_H_, where density for a bound (semi)-ubiquinone was observed in the Q_i_ site (Fig. 3D)^28^. Furthermore, we observed map density features close to the two heme b_L_ groups which likely correspond to ubiquinols bound at the Q_o_ sites (Fig. 3D and E). The distances between the Q_o_ site, heme b_L_ and heme b_H_ within one CIII monomer are consistent with those observed in mammalian CIII_2_ (SI Fig. 5A), with the two heme b_L_ molecules in COB being bridged by a non-canonical UQ (SI Fig. 5B and supplementary text). We detected density for two copies of the flexible UQCRFS1 head domain (Extended Data Fig. 11), which contrasts with a recent work that found only the head domain proximal to the CI quinone tunnel to display flexibility, whereas the distal domain at the CI interface was proposed to be nonfunctional in electron transport^7^. Using focused 3D classification for the distal UQCRFS1 head domain (Extended Data Fig. 11B), we then identified two classes likely representing the extremes of the head domain movement from the B state where the Fe_2_S_2_ cluster is distanced from heme c_1_ to the C state where the Fe_2_S_2_ cluster is closest (Fig. 3F,G, Extended Data Fig. 11C,D)^29, 30^. This movement of the UQCRFS1 head domain is coupled to conformational changes in the unidentified UQCRTT3 protein, thus suggesting a potential role for this subunit in regulation of CIII_2_ activity (Extended Data Fig. 11D). Thus, our observation of the distal UQCRFS1 head domain flexibility, together with the heme b_L_-wedged UQ, suggests that functional symmetry is maintained in the ciliate CIII_2_ despite deviation from the structural symmetry.

To investigate if the membrane-bending capacity of the supercomplex is biologically relevant, we performed electron cryo-tomography of isolated mitochondrial membranes. Cryo-tomograms revealed ∼40-nm tubular cristae densely packed with helical ATP synthase rows and supercomplexes, identified by the conspicuous CI matrix arm (Fig. 4A). To elucidate the supercomplex architecture *in situ*, we performed subtomogram averaging and obtained a map at 28 Å resolution (Extended Data Fig. 1E). The subtomogram average confirmed the presence of the supercomplex, which fits our atomic model (Extended Data Fig. 1F). The appearance of a tubular membrane density in the subtomogram average suggests that the supercomplex adopts a preferred orientation, with its CI-IV_2_ interface approximately aligned with the long axis of the tube (Fig. 4B). Furthermore, the curved membrane region of the supercomplex subtends an angle of ∼130°, indicating that it contributes to the tubular shape of the cristae.

**Figure 4:**
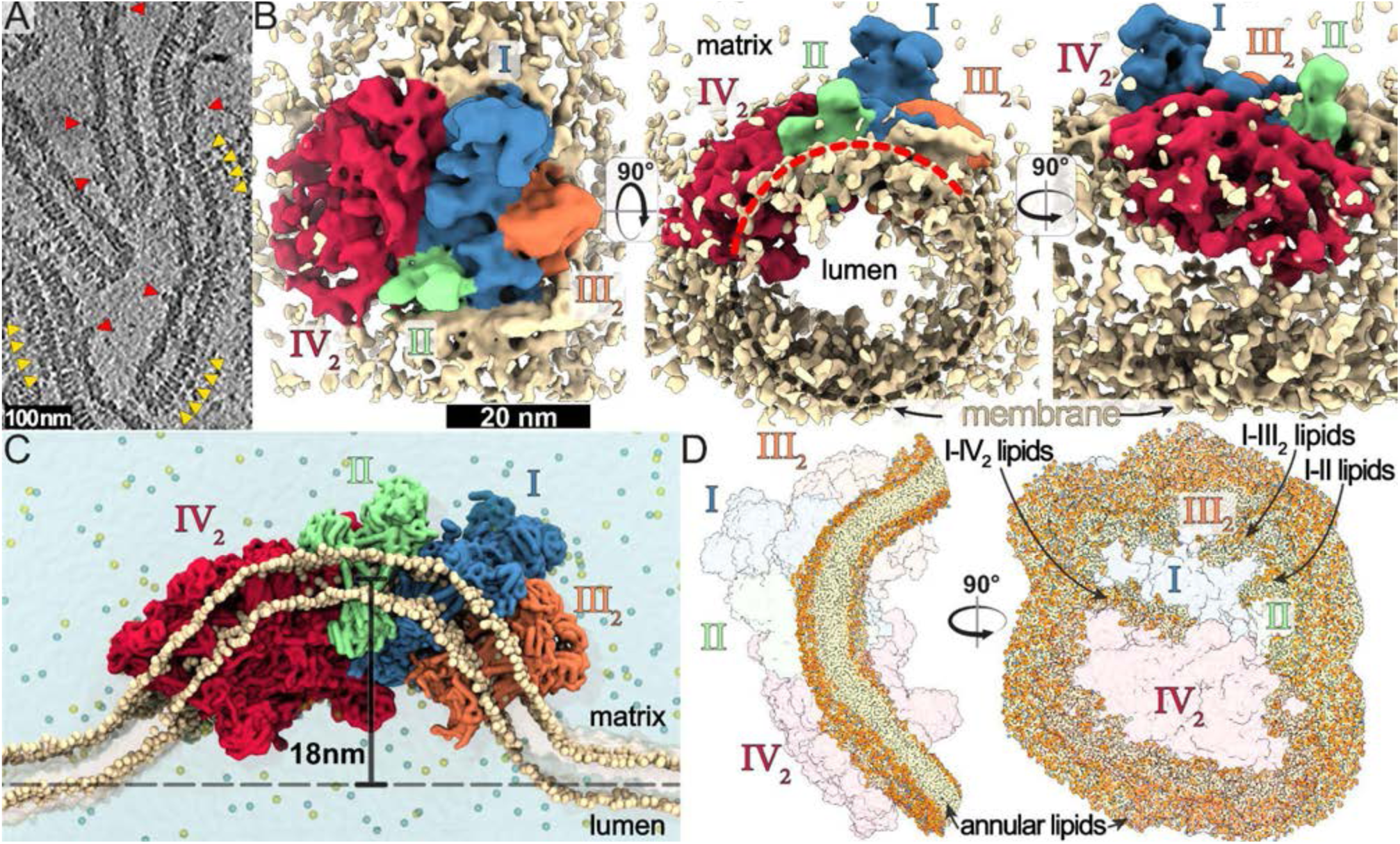
I*n-situ* structure and molecular dynamics of the I-II-III_2_-IV_2_ supercomplex indicate a membrane bending function. **(A)** Cryo-tomographic slice of tubular cristae with ∼40-nm diameter. ATP synthase and supercomplex marked with yellow and red arrowheads. **(B)** Subtomogram average of the I-II-III_2_-IV_2_ supercomplex revealing a preferred orientation in tubular membranes and an arc-shaped structure subtending ∼130° (red dashes). **(C)** Coarse-grained MD simulation showing that the arched membrane region of the supercomplex generates a significant membrane curvature, resulting in 18-nm local displacement of the membrane from the bilayer plane. **(D)** MD simulation reveals the curved structure of the annular lipid shell surrounding the supercomplex, as well as lipid-filled subcomplex interfaces.

To elucidate the membrane-shaping activity of the supercomplex, we performed coarse-grained molecular dynamics simulations. When placed into a planar lipid bilayer, the supercomplex induces a curved membrane topology, displacing the membrane by 18 nm from the original plane (Fig. 4C, SI Video 1). Furthermore, the annular lipid shell surrounding the complex in the equilibrated system displays a highly curved architecture, supportive of an active role in membrane curvature induction (Fig. 4D, SI Video 2). Additionally, we observed lipid pockets in the transmembrane interfaces between subcomplexes, which suggests that their maintenance is crucial for the supercomplex integrity (Extended Data Fig. 2A,B).

Our results indicate a cristae-shaping mechanism involving both the respiratory supercomplex and the ATP synthase to produce membrane tubulation. It serves the function of confining a narrow cristae diameter of around 40 nm, which allows tight cristae packing, thereby increasing the surface area of the bioenergetic membrane. thus favouring ATP synthesis. This membrane- shaping organization of the respiratory supercomplex is markedly different from the mammalian homolog, which resides in the flat crista regions, thereby generating a spatial segregation from ATP synthase^31^. Furthermore, the observed colocalization of the four respiratory complexes would contribute to a directional proton gradient inside the cristae. Because the crista lumen displays the highest membrane potential, with every crista representing an independent functional compartment^32^, the restriction of the cristae diameter likely serves to minimize the lumenal compartment, thereby ensuring that proton translocation results in an increased local membrane potential and ultimately favouring ATP synthesis.

## Methods

### Purification of *T. thermophila* supercomplex

*T. thermophila* cells were grown at the temperature of 36 °C and harvested as previously described^11^, and cell pellets were resuspended in homogenization buffer (20 mM Hepes/KOH pH 7.5, 350 mM D-mannitol, 5 mM EDTA, 1x protease-inhibitor tablet) and lysed in a Dounce homogenizer on ice. Intact mitochondria were isolated first by differential centrifugation of the lysate and finally on a discontinuous sucrose gradient with 15%, 23%, 32% and 60% w/v sucrose in buffer SEM (20 mM Hepes/KOH pH 7.5, 250 mM sucrose, 1 mM EDTA) at 14,1371 xg for 60 min and 4°C in an SW28 rotor. Intact mitochondria sedimented to the interface between 32% and 60% sucrose and were collected from the gradient, snap-frozen in liquid nitrogen and stored at -80°C. The isolated mitochondria were lysed in buffer A (25 mM Hepes/KOH pH 7.5, 25 mM KCl, 5 mM MgCl_2_, 4% w/v digitonin) for one hour on ice. This procedure was previously confirmed as a gentle solubilization method. Following mitochondrial membrane solubilization, cleared lysate was placed on a sucrose cushion (25 mM Hepes/KOH pH 7.5, 25 mM KCl, 5 mM MgCl_2,_ 0.1% w/v digitonin, 30% w/v sucrose) in Ti70 tubes and centrifuged at 164,685 xg for 3 h. The pellet was gently washed and finally resuspended in buffer D (25 mM Hepes/KOH pH 7.5, 25 mM KCl, 5 mM MgCl_2_, 0.1% w/v digitonin). Prior to loading sample material on a size exclusion chromatography column, larger aggregates were pelleted at 30000 xg for 20 min and 4°C. Cleared sample was loaded on a Superose 6 Increase 3.2/300 column equilibrated in buffer D, collecting elution fractions of 100 µL throughout the run. Peak fractions were right away used for cryo-grid preparation.

### UV–visible difference spectroscopy

The sample obtained from the sucrose cushion step was analyzed for heme content and supercomplex composition using UV–visible difference spectroscopy and CN-PAGE combined with in-gel activity assays. UV–visible difference spectra were recorded between 390 and 675 nm using a home-built spectrophotometer. Protein samples were diluted as necessary in 50 mM HEPES, 0.1% digitonin and pH 8.0. Spectra were measured from sodium dithionite-reduced *minus* air-oxidized spectra. When multiple absorption bands overlapped, spectra were deconvoluted using the peak analysis function in OriginPro 2015 (OriginLab Corporation, Northampton, MA, USA).

### Gel electrophoresis

NativePAGE™ 3 to 12%, Bis-Tris, 1.0 mm, Mini Protein Gel (Invitrogen) pre-cast gels were used for Clear Native (CN) PAGE. The gels were loaded with a protein ladder (NativeMark™, Invitrogen) and four identical sample lanes where the protein samples have been mixed with NativePAGE™ Sample Buffer (final concentration, 50 mM BisTris, 6 N HCl, 50 mM NaCl, 10% w/v glycerol, 0.001% Ponceau S, pH 7.2) as per the manufacturer’s instruction.

Electrophoresis was conducted at 4°C, first at 150 V for 30 min with NativePage Light Blue Cathode buffer (50 mM BisTris, 50 mM Tricine, pH 6.8, 0.002% Coomassie G-250) and then at 250 V for 150 min with NativePAGE Anode Buffer (50 mM BisTris, 50 mM Tricine, pH 6.8). In-gel activity assays were performed following published protocols^33^. In brief, each sample lane from CN-PAGE was incubated with an aqueous solution to reveal (i) protein bands (0.02% Coomassie G-250, overnight) or the presence of active (ii) CI (2 mM Tris pH 7.4, 2.5 mg/mL nitrotetrazolium blue chloride (NBT), 0.1 mg/mL NADH, 15 min), (iii) CII (5 mM Tris pH 7.4, 2.5 mg/mL NBT, 84 mM succinic acid, 0.2 mM phenazine methosulfate, 30-40 min) and (iv) CIV (0.05 mM KPi pH 7.4, 0.5 mg/mL 3,3′-diaminobenzidine (DAB), 1 mg/mL cytochrome *c* from *Saccharomyces cerevisiae*, overnight). The reactions were stopped by incubation in 10 % (v/v) acetic acid, followed by multiple exchanges of water. The gel shown is representative of two experiments from two separate supercomplex preparations.

### Cryo-EM sample preparation and data collection

Supercomplex eluted at a concentration of approximately 10 mg/mL. Aliquots of the peak fraction were diluted in buffer D to 0.75 mg/mL before applied to cryo-grids. Quantifoil R2/2- 300 grids floated with a home-made 3 nm amorphous carbon layer were glow-discharged immediately before applying a 3 uL sample. Grids were vitrified using liquid ethane cooled by liquid nitrogen in a Vitrobot Mark IV, with 30 seconds wait time before blotting grids for 3 seconds at blot force 0. Micrographs were collected on a Titan Krios (ThermoFisher Scientific) operated at 300 kV at a nominal magnification of 165 kx (0.83 Å/pixel) with a Quantum K2 camera (Gatan) using a slit width of 20 eV. With an objective lens aperture of 70 µm, images were collected with an exposure rate of 4.26 electrons/pixel/second with 5 seconds exposure fractionated into 20 frames. A total of 26,063 movies were collected.

### Cryo-EM data processing

Motion correction was performed in the internal implementation of RELION-3.1^34^, followed by CTF estimation by CTFFIND 4. Initial rounds of particle picking and 2D classification, followed by ab-initio reconstruction, 3D classification and preliminary refinement of the supercomplex.

Template-based particle picking in RELION was then used to pick and extract 1,664,103 particles. 2D classification and 3D heterogeneous refinement steps in cryoSPARC v.2^35^ were then used to separate supercomplex particles from copurified ATP synthase, resulting in a final 138,746 supercomplex particles used for subsequent refinement. Following a consensus refinement in cryoSPARC, per-particle CTF refinement and bayesian polishing were performed in RELION-3.1. For final refinements in cryoSPARC, particles were downsampled from a 724- pixel box to 480 pixels, resulting in a pixel size of 1.25 Å/px. Masked refinements of the respective supercomplex subregions resulted in map resolutions of 2.9 Å for the entire supercomplex, 2.8 Å for complex-I, 3.0 Å for complex-II, 2.8 Å for complex-III and 2.6 Å for complex-IV_2_. Reported map resolutions are according to gold standard Fourier Shell Correlation (FSC) using the 0.143-criterion. To assess flexibility of the Rieske subunit wedged in the CI-III_2_ interface, we performed focused 3D-classification in RELION-3.1 using pre-aligned particles with a mask on the extended area around the headgroup of the Rieske subunit. Classification into 10 classes resulted in maps confirming flexibility of the structural element, with two classes corresponding closely to the previously reported b- and c-states (Fig. 3F and G).

### Electron cryo-tomography and subtomogram averaging

Crude mitochondrial pellets were resuspended in an equal volume of buffer containing 20 mM HEPES-KOH pH 7.4, 2 mM EDTA, 250 mM sucrose and mixed in a 1:1 ratio with 5-nm colloidal gold solution (Sigma Aldrich) and vitrified as described above on glow-discharged Quantifoil R2/2 Au 200 mesh grids. Tilt series were acquired on a Titan Krios operated at 300 kV with a K3 camera (slit width 20 eV) using serialEM or the EPU software (Thermo Fisher Scientific). Mitochondrial membranes were imaged at a nominal magnification of 42 kx (2.11 Å/pixel) and an exposure rate of 19.5 electrons/pixel/s with a 3 electron/Å^2^ exposure per tilt fractionated into five frames with tilt series acquired using the exposure-symmetric scheme^36^ to ±60° tilt and a 3° tilt increment. Following motion correction in motionCor2, tomographic reconstruction from tilt series was performed in IMOD^37^ using phaseflipping and a binning factor 2. Tomograms were contrast enhanced using nonlinear anisotropic diffusion filtering to facilitate manual particle picking of supercomplex particles based on the matrix arm of complex-I. Subtomogram averaging was performed in PEET^38^. Initial references were generated from the data by averaging after rotating subvolumes into a common orientation with respect to the membrane based on manually assigned vectors. Following initial rounds of averaging to generate a suitable reference, data was manually split into half-sets and refined independently, following lowpass filtering to 50 Å. Averaging of 360 particles from 12 tomograms resulted in a 28-Å subtomogram average.

### Model building and refinement

Manual model building was performed in *Coot*^39^, and new subunits identified directly for the cryo-EM map. For identified canonical subunits, homology models were generated using SWISS modeler. Bound cardiolipins were unambiguously identified from their head group density.

Other natively bound lipids were tentatively modelled as phosphatidylcholine, phosphatidylethanolamine or phosphatidic acid based on head group densities. Real-space refinement of atomic models was performed in PHENIX using secondary structure restraints^40^. Atomic model statistics were calculated using MolProbity^41^.

Given the mild solubilization conditions we used, for CIII_2_ cryo-EM map showed density located on the pseudo-C_2_ symmetry axis between the two COB heme b_L_ molecules displaying planar map features consistent with the quinone moiety of ubiquinone (UQ). Interestingly, the density clearly indicates that UQ can bind in two orientations, related by the symmetry rotation of the dimer. In either of the two orientations, the quinone moiety is positioned close to a heme b_L_, where potentially it could accept electrons for transfer across the dimer axis. In the recent amphipol CIII_2_ structure^7^, the isoprenoid tail of UQ was modeled in the equivalent position, however, planar density for the quinone was missing. This orientation-equivalent binding of UQ between the two COB heme b_L_ molecules, together with the B- and C-state Rieske conformations, suggest a maintained functional symmetry of ciliate CIII_2_ within the supercomplex.

In the CI, we identified 49 canonical subunits and 21 subunits that we assign as phylum-specific. In each CIV monomer, we identified 11 subunits homologous to mammalian CIV (COX1, 2, 3a, 3b, 5B, 6A, 6B, 6C, 7A, 7C, NDUFA4) and 42 ciliate-specific subunits, most of which are peripherally associated around the mitochondrial protein core. Three of the mammalian subunits missing in *T. thermophila* CIV (COX4, COX7B, and COX8) are at the interface where two mitochondrial carriers are bound. The mitochondrially encoded core subunit COX3 is split into two fragments. Most of the TM helices are contributed by the C-terminal COX3b, which is encoded by the mitochondrial *ymf68* gene. The newly annotated Ymf68 is structurally conserved, apart from the missing helix (H1), which is structurally replaced by Ymf67. We therefore assign *ymf67* and *ymf68* of the ciliate mitochondrial genome as separately encoding the COX3a/b subunit fragments. On the *T. tetrahymena* mitochondrial genome, *ymf67* and *ymf68* genes are located on the same strand, but separated by the gene for tRNATrp, suggesting that a transposition event may have led to the fragmentation of the original COX3 gene. tRNA genes are known to be among the most motile elements in metazoan mtDNA. Both COX3a/b fragments have evolved substantial subunit extensions threading through the augmented CIV monomer unit to recruit lineage-specific subunits and mediate supercomplex assembly.

In the CIV dimer, the dimer interface of 17,000-Å^2^ is dominated by 16 species-specific subunits. Furthermore, when aligned on the CIV core, a comparison of the mammalian and ciliate structures reveals that the two dimers display markedly different architectures, dimer axes and distances between COX1 cores. This suggests that the ciliate CIV dimerization likely evolved through the acquisition of lineage-specific subunits and reflects the constraints of the unique tubular membrane environment.

In addition, each CIV monomer complex contains two different Surf1-like proteins, which in human were reported to complement defects causing the Leigh syndrome. In our structure, two Surf1-like proteins are permanently attached to CIV and display similar overall structures, consisting of a lumen-exposed soluble domain and a transmembrane-helix hairpin. The two Surf1 proteins are facing each other, bound on opposite sides of each CIV core.

The presence of subunit extension and accessory subunits in CIV generates a pronounced cavity around the cytochrome *c* binding site. However, overlaying of a *Tetrahymena* cytochrome *c* homology model suggests that the canonical binding site is not obstructed. Cytochrome *c* binding is known to be driven by electrostatic interactions with the CuA domain of COX2, which in mammals forms a negatively charged patch. This structural feature is positively charged in *T. thermophila*, interacting mainly with H1 of cytochrome *c*, which displays a flipped polarity. We conclude that the experimentally observed functional incompatibility of *Tetrahymena* CIV and mammalian cytochrome *c* is not due to divergent architecture, but an inverted surface charge of the binding pocket.

### Molecular dynamics simulations

We performed coarse-grained (CG) molecular dynamics (MD) simulations on the entire *Tetrahymena thermophila* supercomplex structure using Martini3 forcefield^42^ to study the rearrangement of the lipid bilayer around the highly bent protein assembly. Using the *martinize2* (version 2.6) tool, we transformed the atomistic structure into a CG model (atoms clustered into Martini beads)^43^. Using the small molecules database and the existing topologies of phospholipids available in the Martini3^42^ forcefield, we generated the force field parameters for cardiolipin. Cofactors and resolved lipids in the structure were not included in the simulated model system, and only protein was simulated to study the dynamics of lipid molecules around it. First, the CG model of the protein structure was minimized for 100 steps in vacuum to remove possible steric clashes. Then the minimized CG supercomplex was embedded in a large (75 nm x 75 nm) hybrid membrane slab (POPE:POPC:CL in 4:2:1 ratio) using the *insane.py* script^44^. The coarse-grained protein-membrane system was solvated using standard Martini3 water beads and 100 mM Na^+^ and Cl^-^ ions. Starting from this initial position, the simulation system was minimized keeping all beads free; first in double precision to resolve steric clashes between the lipids (maximum 500 steps) and then in regular single precision (maximum 10 000 steps). After minimization, with 4000 kJ mol^-1^ nm^-2^ harmonic constraints on the backbone beads, the system was equilibrated using velocity-rescaling thermostat^45^ and Berendsen barostat^46^ for 10 nanoseconds. During production runs, the 4000 kJ mol^-1^nm^-2^ harmonic constraints on the backbone beads were applied. Velocity-rescaling thermostat^45^ and Parrinello-Rahman barostat^47^ were used for temperature (310 K) and pressure (1 bar) control in the production phase.

Coulombic interactions were treated with the reaction-field algorithm using ε_r_ = 15^48^. The Verlet cutoff scheme was implemented with a Lennard-Jones cutoff of 1.1 nm^49^. The time step of the coarse-grained MD simulations was 20 femtoseconds. Initial simulation replicas showed incomplete or unstable wrapping of the membrane around the protein, so we translated the lipid bilayer patch in z-direction and altered the insertion angle of the supercomplex to find an initial position that allowed the membrane to equilibrate and wrap fully around the protein (Systems T1-T7, simulation lengths 0.9 – 2.8 microseconds, total 9.6 microseconds). After finding the correct insertion of the protein into the membrane, we initiated three independent simulation replicas (Systems P1-P3, simulation lengths 10 microseconds each, total 30 microseconds). The simulations were performed using the Gromacs software (version 2021)^50^.

### Data visualization and analysis

Images were rendered using ChimeraX^51^. To analyze the *T. thermophila* cytochrome-c binding site, the mammalian cytochrome-c bound complex-IV structure (PDB 5iy5) was overlaid with the *T. thermophila* structure. Using AlphaFold2^52^, a *T. thermophila* cytochrome-c structure was predicted and overlaid into both the mammalian structure and *T. thermophila* structures. The composite map of the complete respiratory supercomplex was generated in ChimeraX^51^. This map was only used for visualization, but not for atomic model refinement, where instead a consensus map was used. The buried areas of the CI-CII-CIII_2_-CIV_2_ interfaces and CIV dimer interface was calculated in ChimeraX^51^.

## Data availability statement

The atomic coordinates were deposited in the RCSB Protein Data Bank (PDB) under accession number XXXX. The cryo-EM maps have been deposited in the Electron Microscopy Data Bank (EMDB) under accession number EMD-XXXXX.

## Acknowledgments

We thank SciLifeLab EM facility (funded by the KAW, EPS, and Kempe foundations), EMBO Young Investigator Program, Biological Physics group at the University of Helsinki and Dr. Paulo C. T. Souza for discussions on MD simulations; Swedish Foundation for Strategic Research (FFL15:0325), Ragnar Söderberg Foundation (M44/16), Cancerfonden (2017/1041), European Research Council (ERC-2018-StG-805230), Knut and Alice Wallenberg Foundation (2018.0080), V.S. was supported by Academy of Finland, Sigrid Jusélius Foundation, Jane and Aatos Erkko Foundation, Magnus Ehrnrooth Foundation and the University of Helsinki, A. Ma. was supported by Medical Research Council (CDA MR/M00936X/1, Transition Support MR/T032154/1).

## Author contributions

A.M., R.K.F. and A.A. designed the project. V.T. performed cell culturing and isolation of mitochondria. A.M. and R.K.F. prepared the sample and collected cryo-EM data. A.M., R.K.F. and R.B. processed cryo-EM data and built the model. A.M. performed cryo-ET and subtomogram averaging. O.H. and V.S. performed molecular dynamics simulations. T.G, A.Ma., and A.A. performed biochemical and spectroscopic analyses. A.M., R.K.F. and A.A. wrote the manuscript with contributions from O.H., V.S and A.Ma. All authors contributed to revising the manuscript.

---

**Extended Data Fig. 1.**
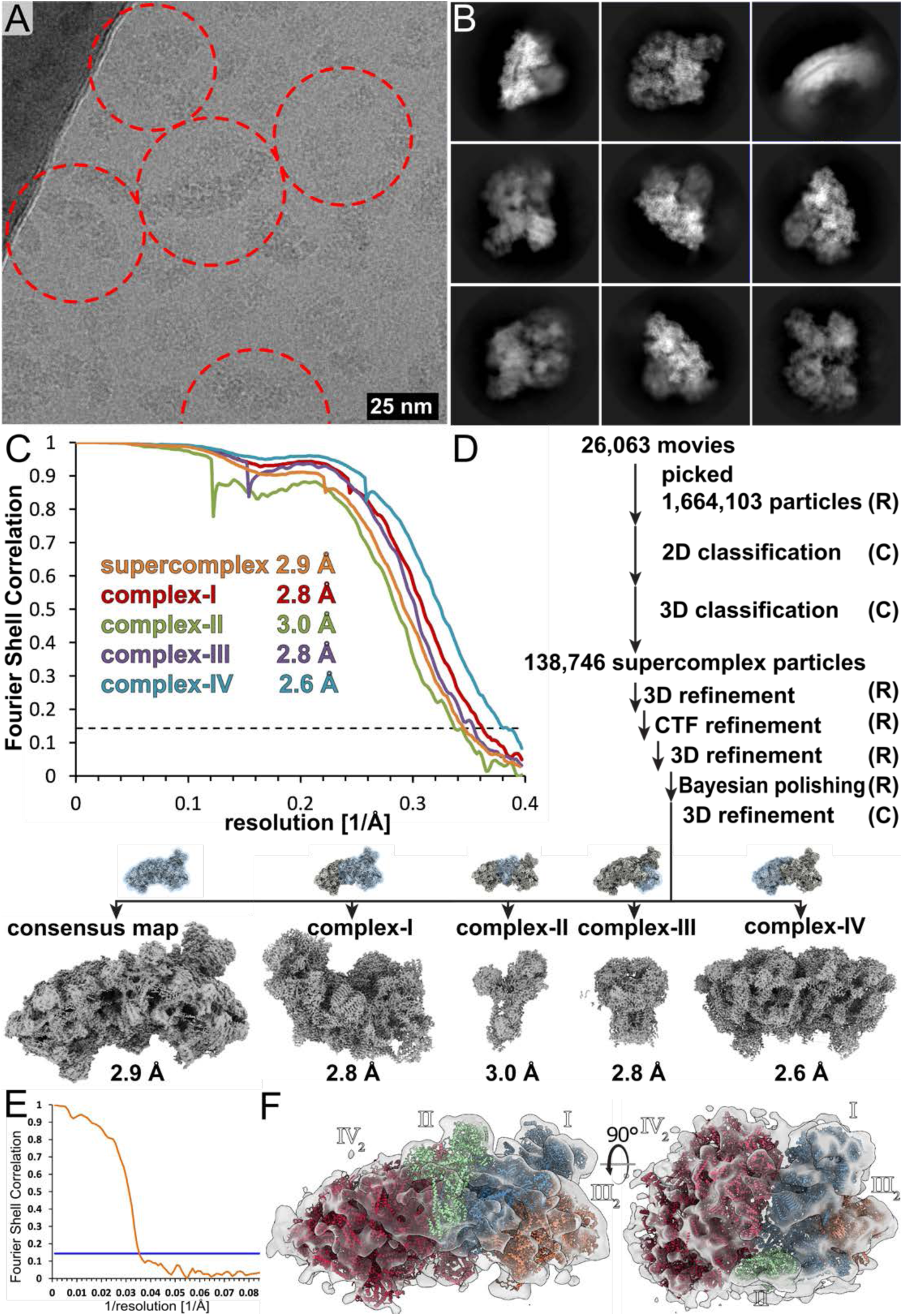
Cryo-EM and subtomogram averaging data processing. **(A)** representative micrograph with supercomplex particles indicated. **(B)** 2D class averages. **(C)** Fourier Shell Correlation of the five final maps according to the 0.143 gold-standard criterion. **(D)** particle processing workflow in RELION-3.1 (R) and cryoSPARC2 (C). **(E)** Fourier Shell correlation of the subtomogram average indicating a resolution of 28 Å. **(F)** The subtomogram average map (transparent grey) agrees well with the atomic model of the I-II-III_2_-IV_2_.

**Extended Data Fig. 2.**
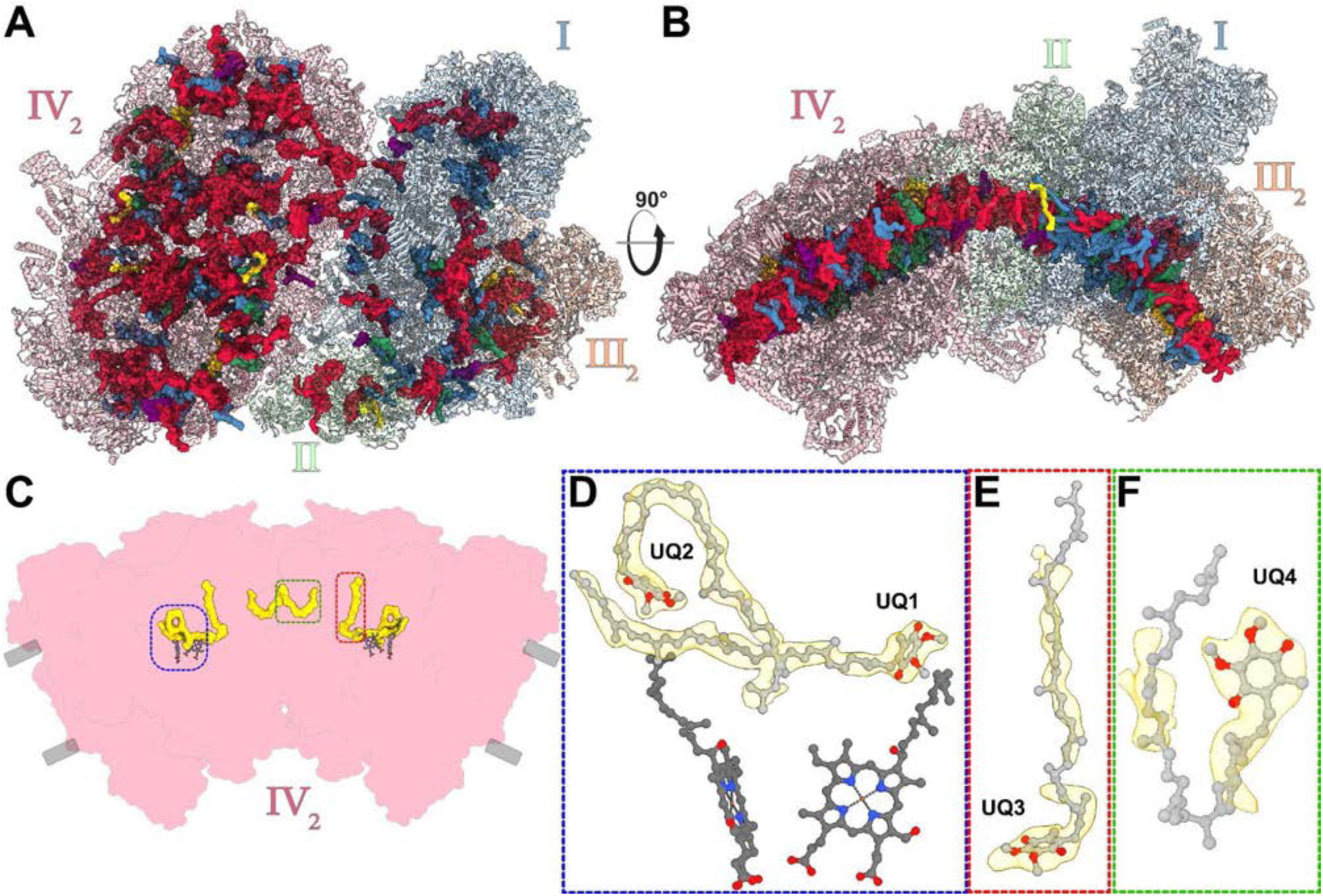
Bound native lipids of the ciliate I-II-III_2_-IV_2_ supercomplex reveal a curved membrane region. Top view **(A)** and side view **(B)** of the supercomplex structure with bound lipids cardiolipin (red), phosphatidylcholine (blue) phosphatidylethanolamine (green), phosphatidic acid (yellow), ubiquinone-8 (purple) spread throughout the membrane region. **(C-F)** Location of the four ubiquinone-8 molecules found in each CIV monomer with closeup views **(D-F)**, including two UQ-8 molecules bound close to the heme centres **(D)**.

**Extended Data Fig. 3.**
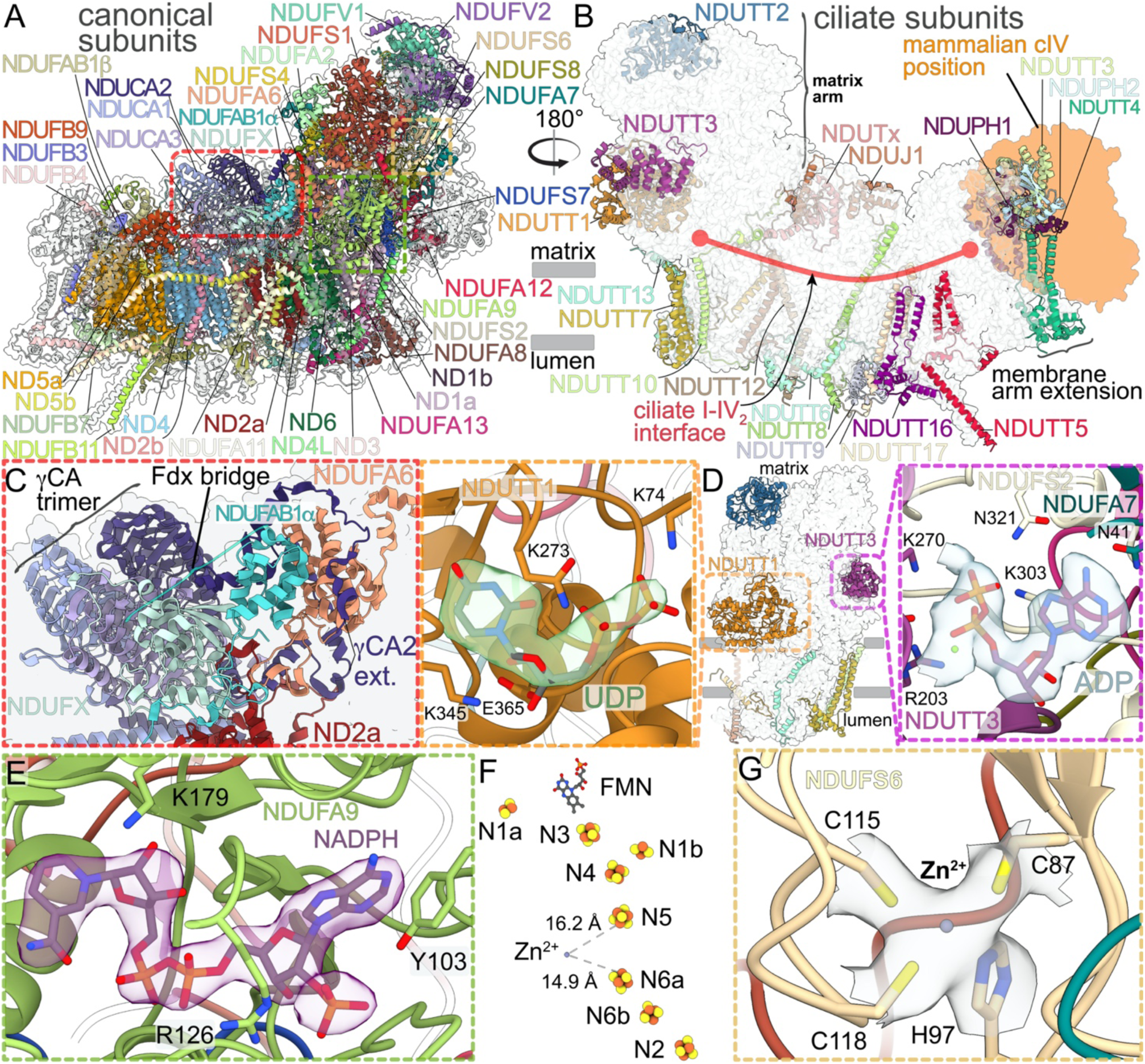
Lineage-specific structural features of CI dictate a divergent supercomplex architecture. The canonical subunits of CI (coloured) include the conserved antiporter-like folds (ND2a/b, ND4, ND5a/b), as well as a γ-carbonic anhydrase trimer. **(B)** The augmented CI architecture occludes the canonical CIV binding site. **(C)** The ferrodoxin bridge of NDUAB1-a and NDUFX connects ND2a of the membrane arm and NDUA6 of the matrix arm. **(D)** The matrix arm contains bound soluble subunits NDUTT1 with a bound UDP and NDUTT3 with a bonus ADP. **(E)** NDUFA9 contains a bound NADPH molecule located close to the CI Q- tunnel. **(F)** Overview of redox active centers in CI matrix arm including the **(G)** zinc ion coordinated by three Cys and one His residue within NDUS6.

**Extended Data Fig. 4.**
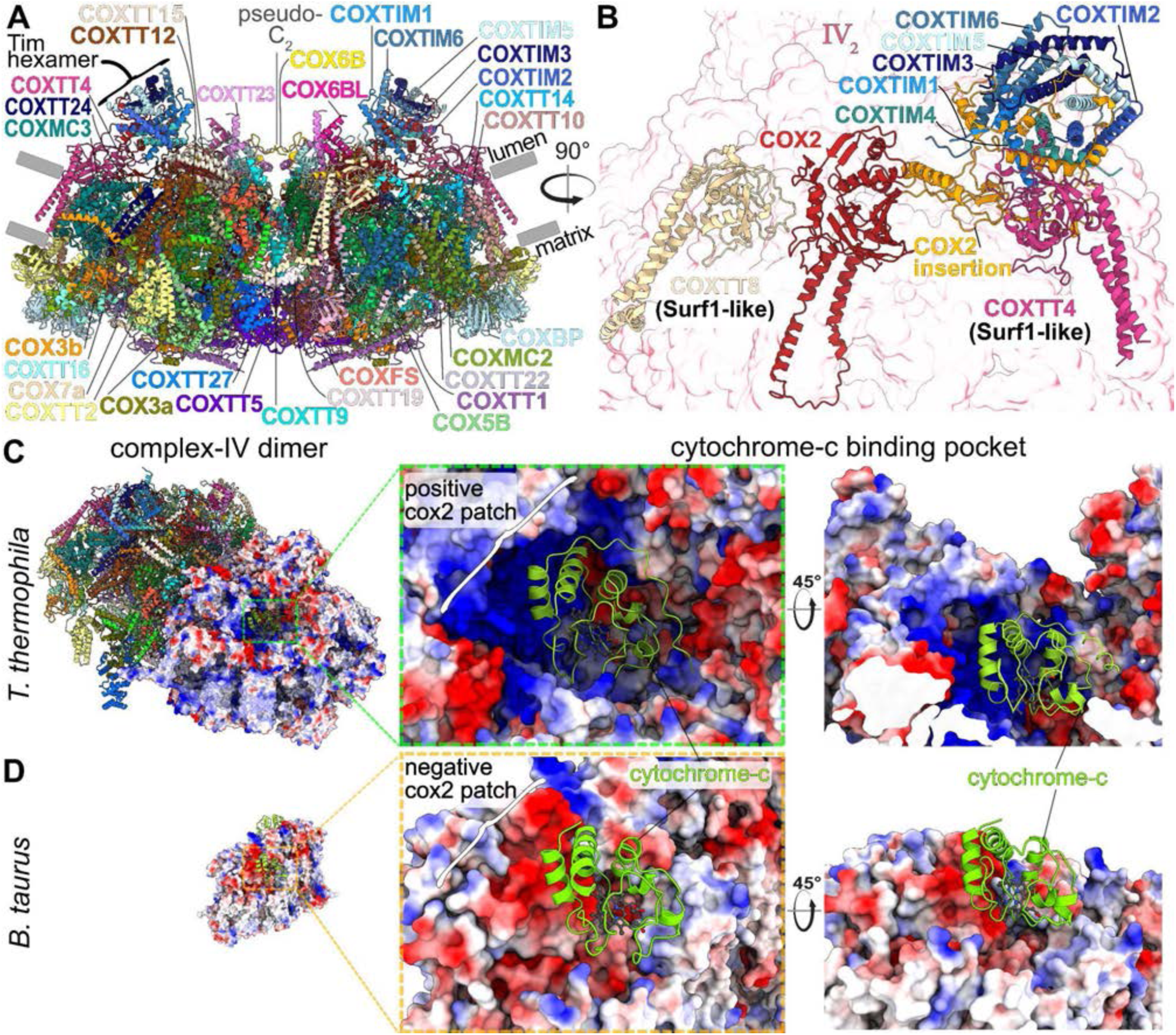
CIV_2_ contains a bound Tim hexamer and a cytochrome*-c* binding site with inverted electrostatic charge. **(A)** sideview of CIV_2_, containing numerous accessory subunits, including a Tim hexamer (B) Closeup view of COX2, which contains an insertion that recruits the Tim hexamer to the CIV dimer. Furthermore, COX2 interacts with the two Surf1-like proteins COXTT8 and COXTT4. **(C-D)** ciliate (C, this study) and mammalian (D, PDB 5IY5) CIV_2_ structures. An overlay of a predicted Tt-cytochrome-c structure fits the cytochrome-c binding crater without clashes, suggesting that the difference in binding affinities is derived from inverted surface charge of the COX2 patch on the binding site.

**Extended Data Fig. 5.**
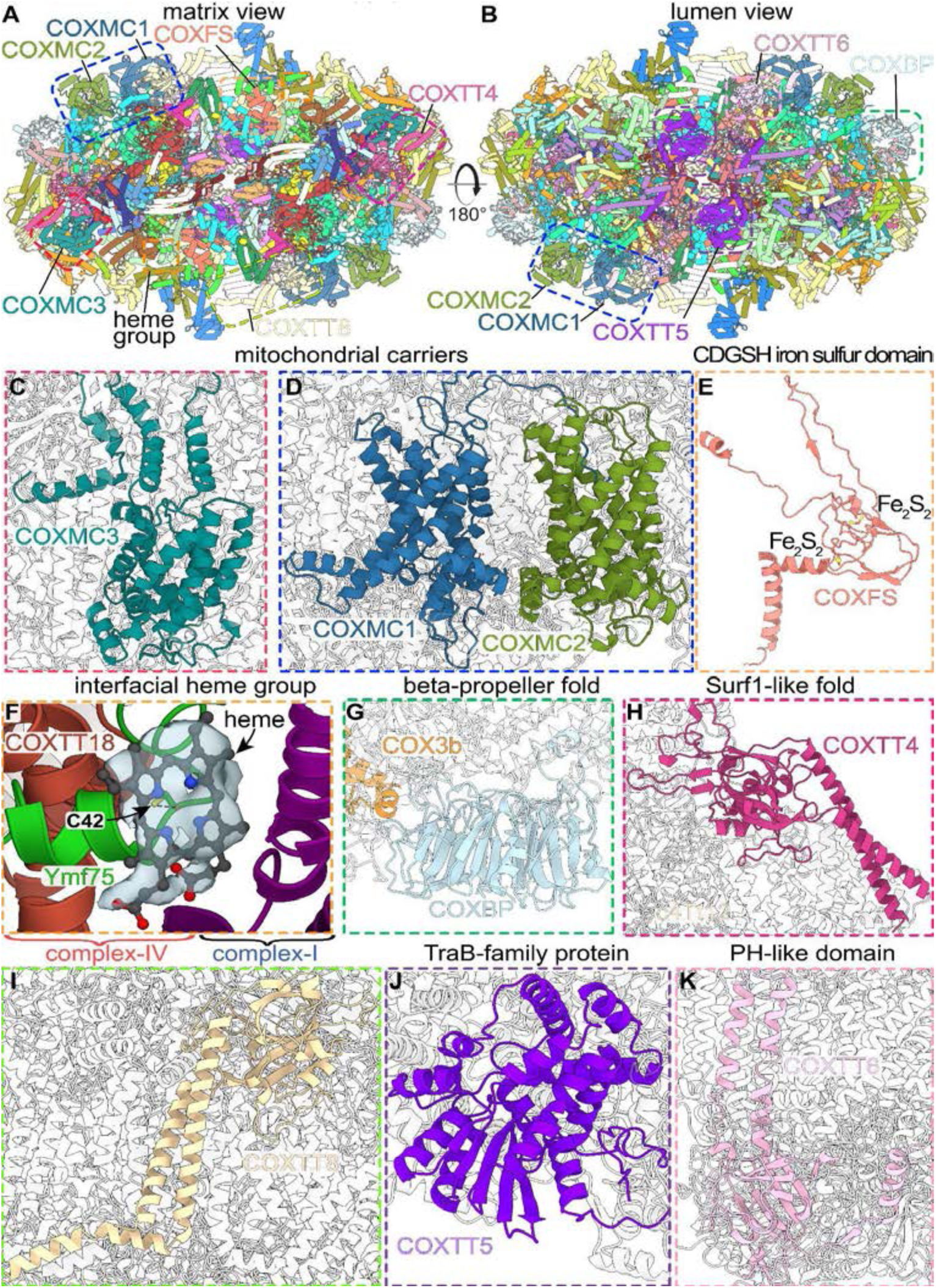
The augmented CIV dimer contains numerous associated compact- fold subunits. **(A,B)** Matrix and lumen view of the CIV dimer, subunit locations of insets are indicated. (**C,D**) Subunits COXMC1,MC2 and MC3 form mitochondrial carrier folds. COXFS is a CDGSH iron sulfur domain **(E).** A similar recruitment of a CDGSH-like protein has been found in the *T. thermophila* mitochondrial ribosome, where mL107 plays a structural role in the large mitochondrial subunit (Tobiasson et al). **(F)** Ymf75 coordinates a noncanonical heme group at the CIV periphery. (**G-K**) Compact folds of accessory subunits include a seven-bladed beta-propeller (G), two Surf1-like proteins (H, I) a TraB-family protein (J) and a Pleckstrin homology (PH) like domain (K).

**Extended Data Fig. 6.**
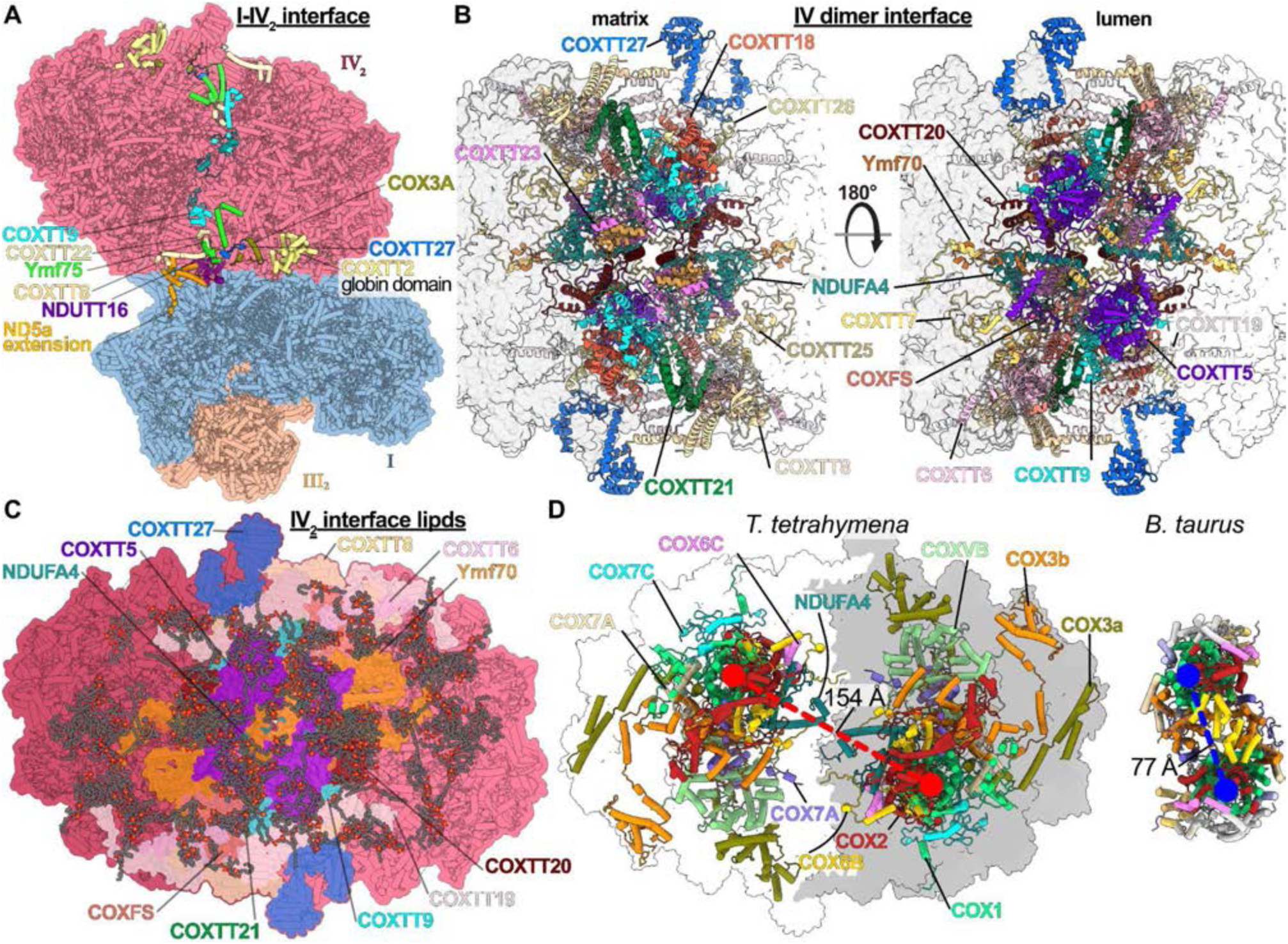
Formation of an augmented CIV dimer enables divergent supercomplex assembly. **(A)** The I-II-III2-IV2 supercomplex displays additional subunits and ordered structural elements (colored cartoons) compared to the amphipol embedded, isolated CIV_2_ dimer (PDB 7W5Z). Additional features of the CI-IV interface include subunit NDUTT16 as well as an entire globin-like domain of COXTT2. **(B)** Matrix and lumenal views of the CIV_2_ interface include NDUFA4. **(C)** The CIV_2_ contains 210 lipids, many of which populate the dimer interface region. **(D)** The *T. thermophila* dimer is augmented compared to the bovine dimer structure and contains the previously unassigned canonical subunit NDUFA4, which is absent in the bovine dimer (right, PDB 3X2Q). Vectors (red, blue) indicate different distances between COX1 centers.

**Extended Data Fig. 7.**
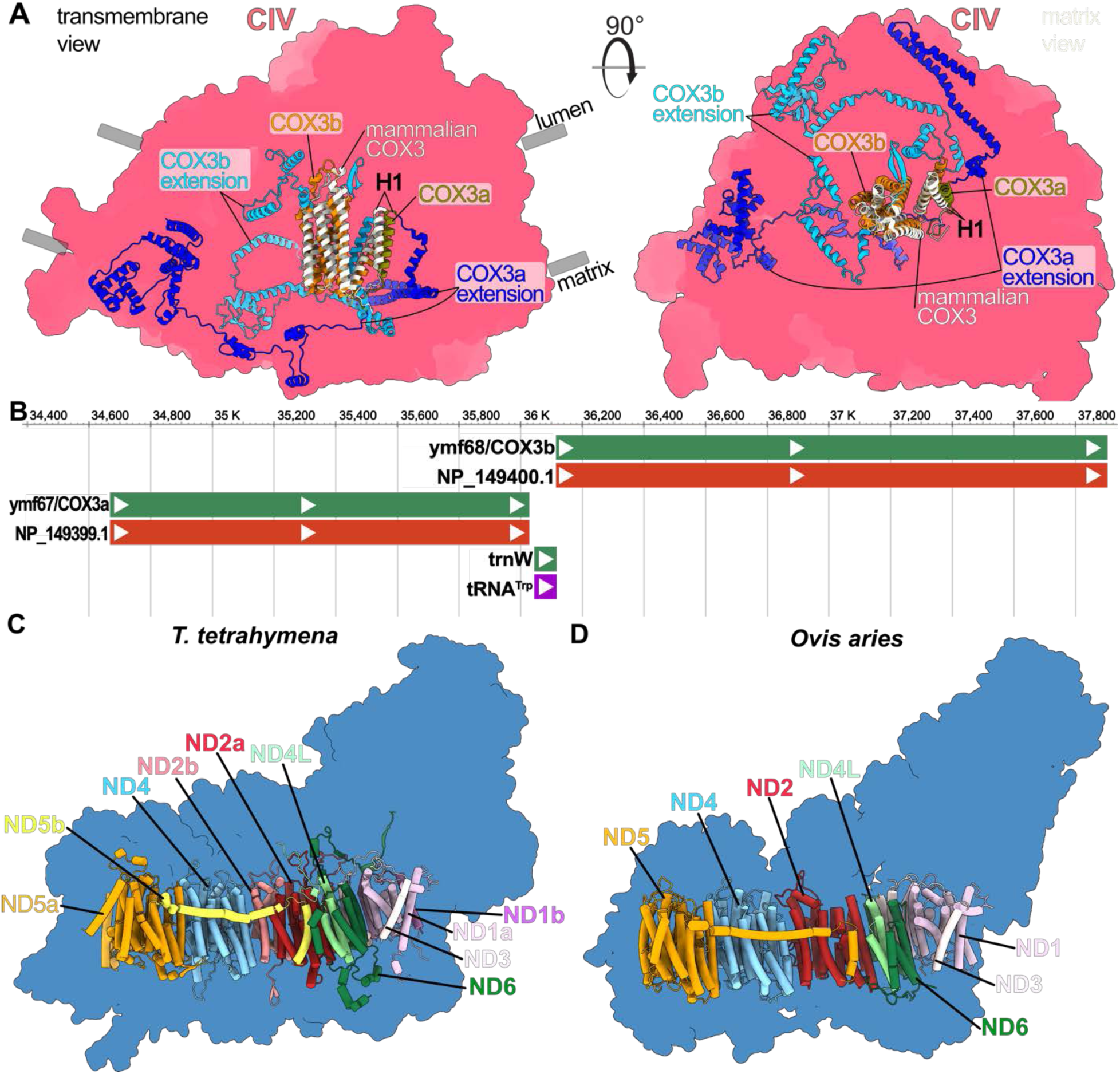
Fragmentation and extension of COX3 and CI proton-pump subunits. **(A)** Overlay of the mammalian COX3 (PDB 5IY5) and the Tt-COX3 (this study) shows that the N-terminal fragment (COX3a), encoded by *ymf67* corresponds to Helix-1 (H1) of the conserved structure. COX3b, encoded by *ymf68* makes up most of the conserved COX3 fold and contains large extensions that contribute to accessory subunit recruitment and supercomplex formation. **(B)** *T. thermophila* mitochondrial genome region showing the insertion of the tRNA-Trp gene in between *ymf67*/COX3a and *ymf68*/COX3b (genes in green, protein/RNA transcripts in red and purple). **(C)** ciliate CI outline (blue) with the antiporter-like subunits shown in color-coded ribbons. The canonical proton pumps ND1, ND2 and ND5 are encoded by split genes, thus resulting in chimeric proteins with a and b chains. **(D)** mammalian CI outline (blue, PDB 5LNK) with subunits shown in ribbons color-coded as in A, showing the single-chain ND1-6 proteins.

**Extended Data Fig. 8.**
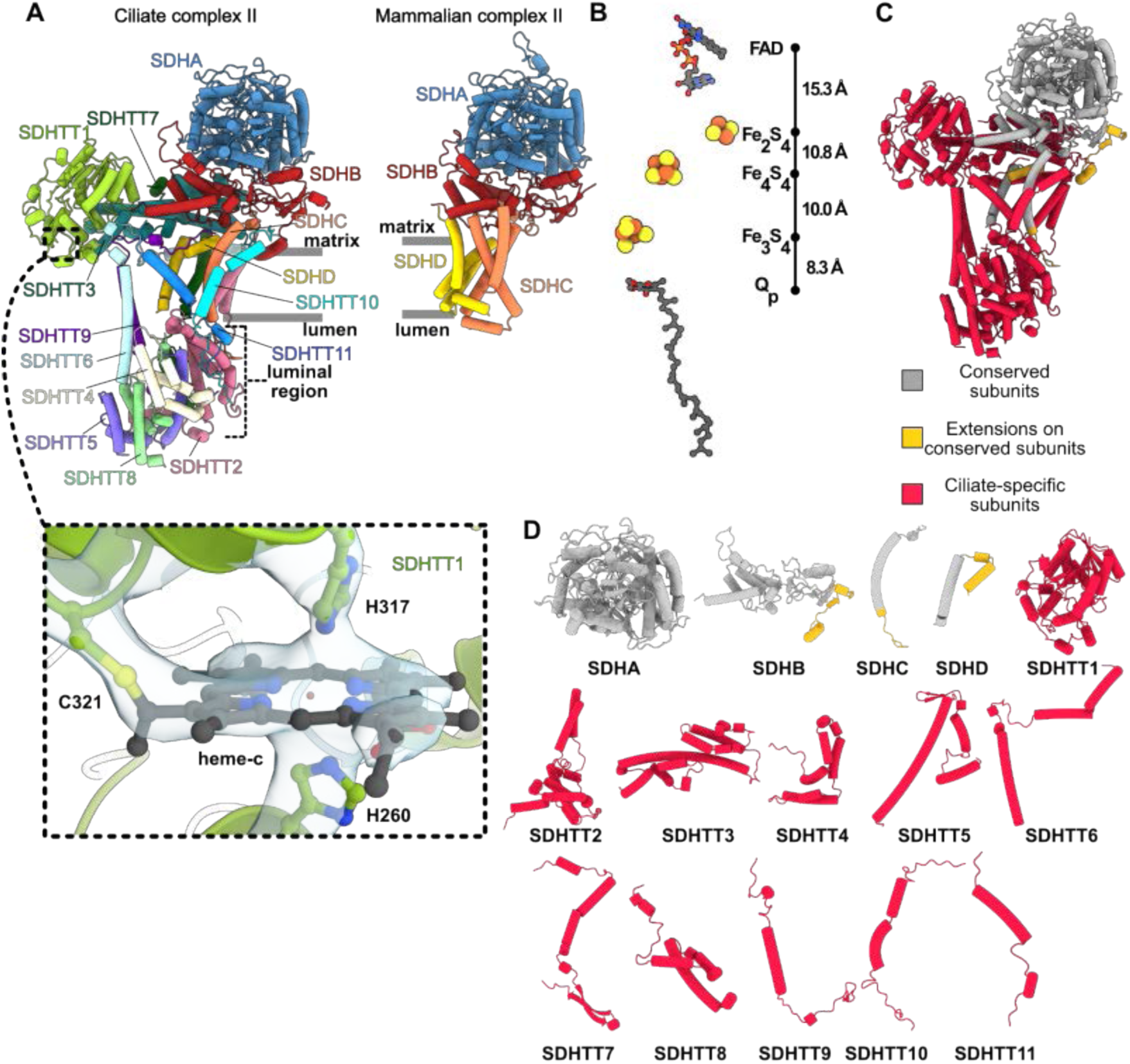
Conservation and divergence of ciliate CII. **(A)** Ciliate CII (left) colored by subunits and shown in comparison to mammalian CII (right, PDB 1ZOY) with similar color scheme for subunits SDHA-D. Evidently, the ciliate SDHC and SDHD subunits are substantially reduced in size and replaced by ciliate-specific subunits. Ciliate CII contains a sizable luminal protein region, whereas mammalian CII contains no equivalent region. Dashed window on SDHTT1 marks the view of a noncanonical cysteine-linked heme C that is coordinated by two axial histidines in SDHTT1E. **(B)** Conserved electron transfer chain in CII.**(C)** CII colored by conserved subunits (grey), extensions to conserved subunits (gold) and ciliate-specific subunits (red). Conservation is assessed as structural homology to mammalian CII (PDB 1ZOY). The ciliate CII shows a high degree of conservation for the four core SDHA-D subunits, with the ciliate-specific subunits essentially making up all the luminal domains, half of the transmembrane domains and half of the matrix domains. **(D)** Individual CII subunits shown as gallery with similar color coding as in C. Subunit naming follows the scheme in (*7*) with SDHTT1 being of largest Mw and SDHTT11 of the smallest molecular weight.

**Extended Data Fig. 9.**
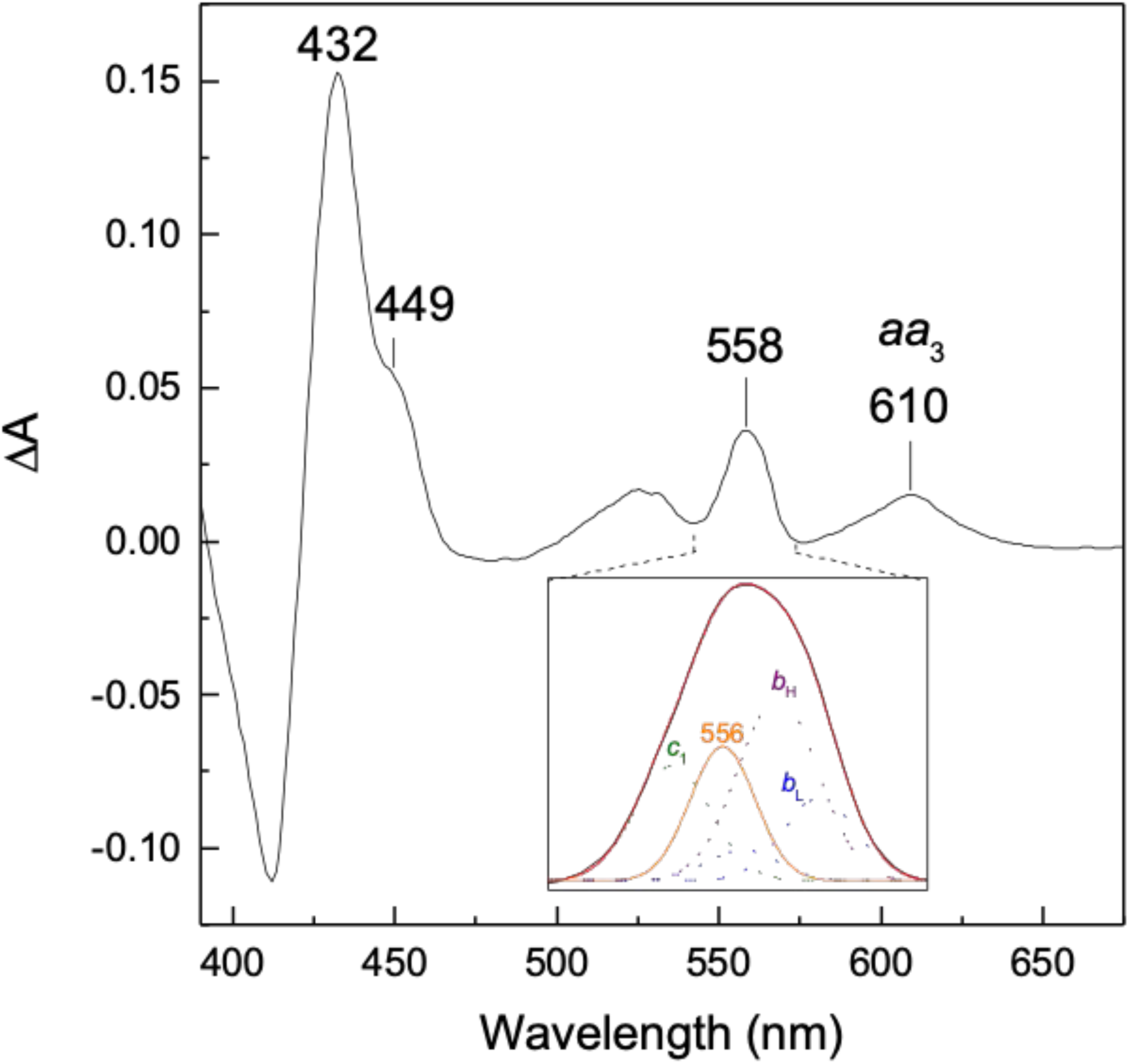
UV-visible redox absorption spectrum of the supercomplex. The dithionite minus air-oxidized difference spectrum was recorded in a 0.3 cm quartz cuvette after twofold dilution of the final sample in 50 mM HEPES, 0.1% digitonin, pH 8.0. Peaks are labeled that correspond to the absorption of the A-type hemes present in CIV (449/610 nm, *aa*_3_) as well as merged absorption bands of other B- and C-type hemes (maxima at 432/558 nm). Inset: Deconvolution of the 558-nm absorption band to highlight the contribution of the B- and C-type hemes of CIII (*b*_H_ at 561 nm, purple; *b*_L_ at 558 and 565 nm, blue; *c*_1_ at 552 nm, green) and the presence of at least another heme-protein with absorption maximum at 556 nm (orange). The spectrum shown is representative of two separate supercomplex preparations.

**Extended Data Fig. 10.**
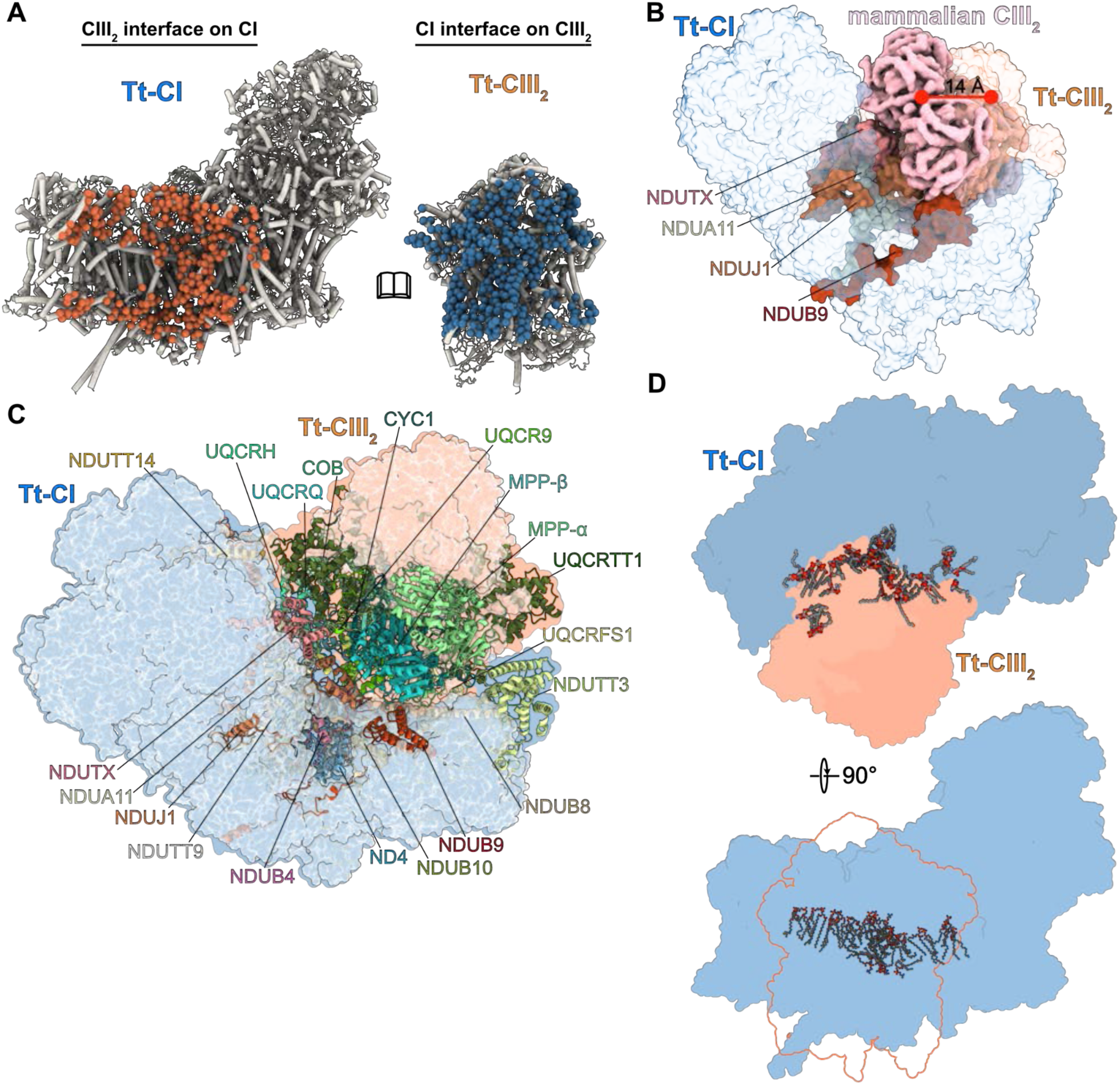
CI-CIII_2_ interface is extensive and involves numerous lipids. **(A)** Buried surfaces in the CI-CIII_2_ interface. Orange spheres show CIII_2_ contacts on CI, blue spheres show CI contact points on CIII_2_. Both sides of the CIII dimer interact with CI resulting in a >9,000 Å^2^ interface. **(B)** Matrix view shows that four Tt-CI subunits would clash with mammalian CIII_2_ in canonical position, thus Tt-CIII_2_ is displaced 14 Å from Tt-CI. **(C)** CI outline (light blue surface) and CIII_2_ outline (light orange surface) highlights the docked position of CIII_2_ on the concave side of the bent CI membrane arm. All subunits that participate in CI-CIII_2_ interface interactions are shown in ribbons and colored as in Fig. 2A,B and Fig. S6A. Non-interacting subunits of CI and CIII_2_ are shown in transparent, light grey surface. **(D)** Orthogonal views of bound native lipids in the I-III_2_ interface. Lipids are distributed in both matrix and luminal leaflets.

**Extended Data Fig. 11.**
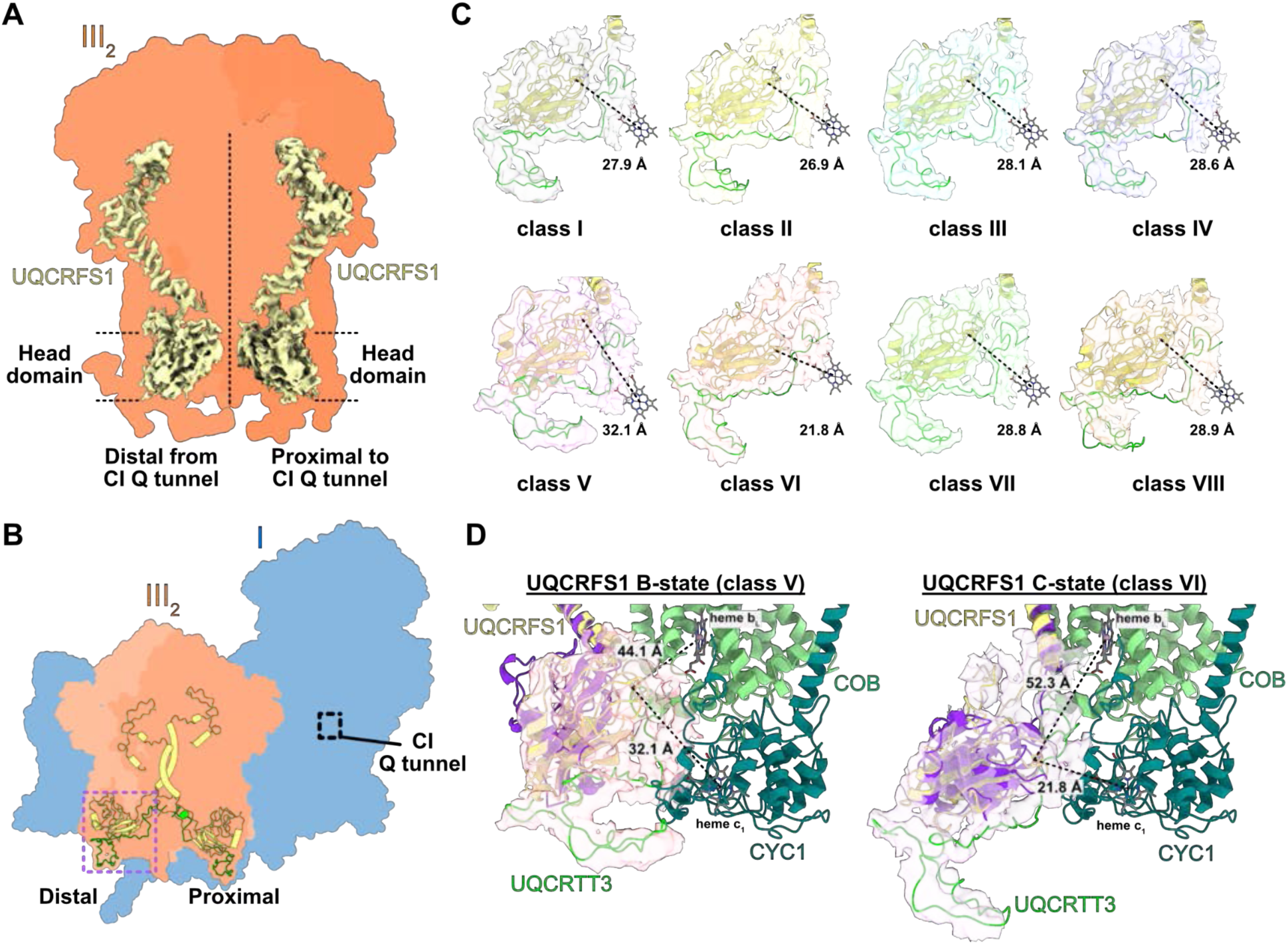
Conformational flexibility in distal UQCRFS1 head domain. **(A)** cryo-EM map density carved around the UQCRFS1 proteins in CIII_2_. Head domain map density for both copies show similar features of inherent flexibility. **(B)** location of distal and proximal UQCRFS1 subunits (yellow ribbon) with respect to CI quinone tunnel (black dashed window). The purple box highlights the region targeted in focused 3D classification, which is the distal UQCRFS1 head domain. UQCRTT3 protein is shown in green ribbon. **(C)** Gallery display of eight 3D classes I-VIII. The distal UQCRFS1 protein and UQCRTT3 were fitted to each 3D class density map and distance measured from Fe_2_S_2_ cluster to heme c_1_ fixed in consensus refined 3D map. Only classes V and VI display markedly different distances. **(D)** Comparison of B-state class V (left) and C-state class IV (right). B-state UQCRFS1 superposes well with stigmatellin-induced B-state UQCRFS1 from chicken CIII_2_ (purple ribbon, PDB 3BCC). C-state UQCRFS1 superposes almost perfectly with C-state UQCRFS1 from chicken CIII_2_ (purple ribbon, PDB 1BCC). Distance measures are shown for ciliate UQCRFS1 Fe_2_S_2_ to heme c_1_ and b_L_.

## SUPPLEMENTARY INFORMATION

### Table of contents

SI Figure 1. Conservation and divergence of ciliate CI

SI Figure 2. CN-PAGE gel showing active CI, CII and CIV in the final supercomplex sample

SI Figure 3. Conservation and divergence of ciliate CIV

SI Figure 4. Conservation and divergence of ciliate CIII

SI Figure 5. Ciliate CIII_2_ heme group distances and wedged ubiquinone

SI Table 1. Data collection and model statistics

SI Table 2. List of proteins and comments

SI Video 1. Coarse-grained molecular dynamics simulation of the *T. thermophila* supercomplex.

SI Video 2. Annular lipid shell of the *T. thermophila* supercomplex.

**Supplementary Information Figure 1.**
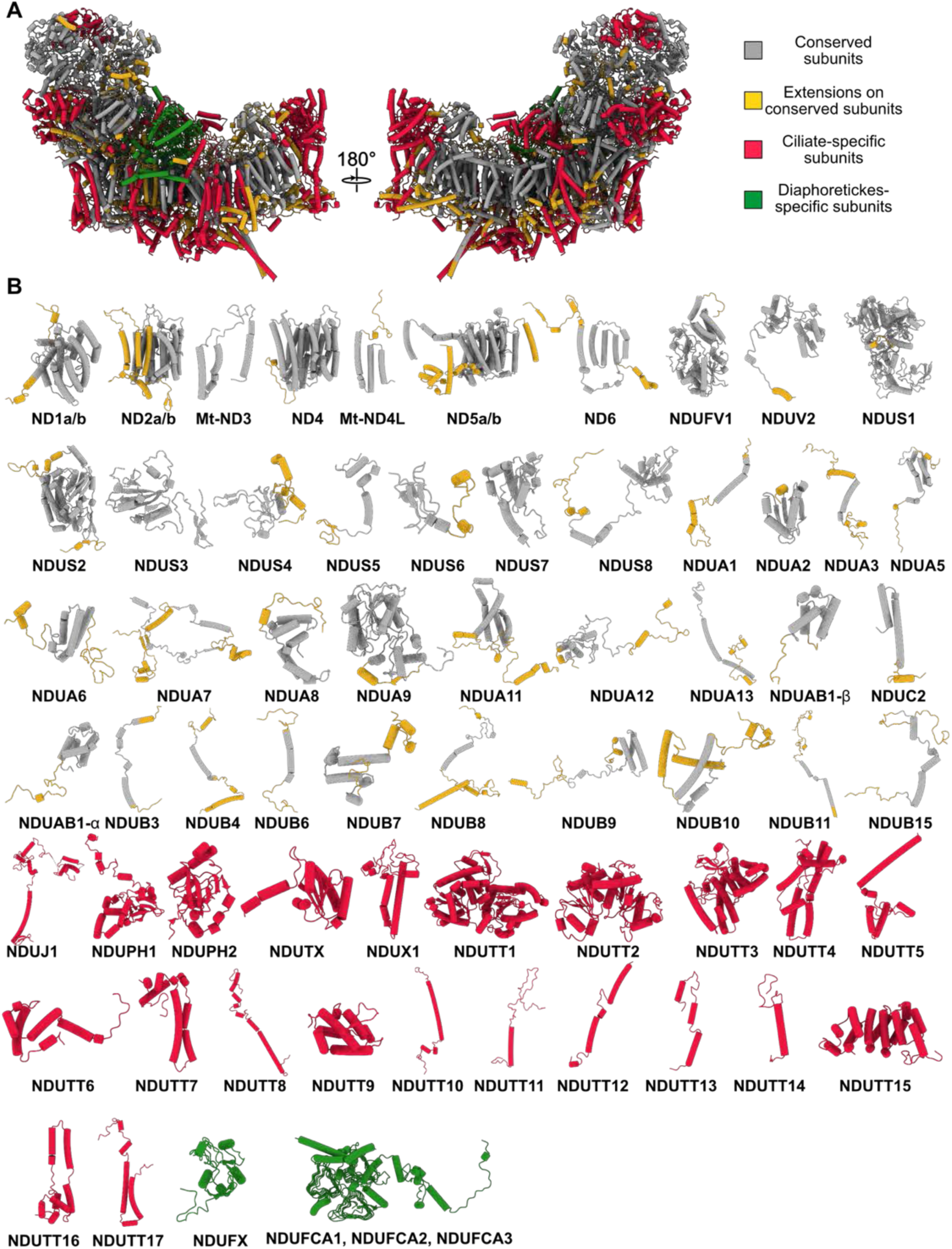
Conservation and divergence of ciliate CI. **(A)** CI colored by conserved subunits (grey), extensions to conserved subunits (gold), ciliate-specific subunits (red) and Diaphoretickes-conserved subunits (green). Conservation is assessed as structural homology to mammalian CI (PDB 5LNK). Extensions and ciliate-specific subunits primarily locate to peripheral regions of the matrix arm and membrane domain. **(B)** Individual CI subunits shown as gallery with similar color coding as in A. Subunit naming as in ref (*7*).

**Supplementary Information Figure 2.**
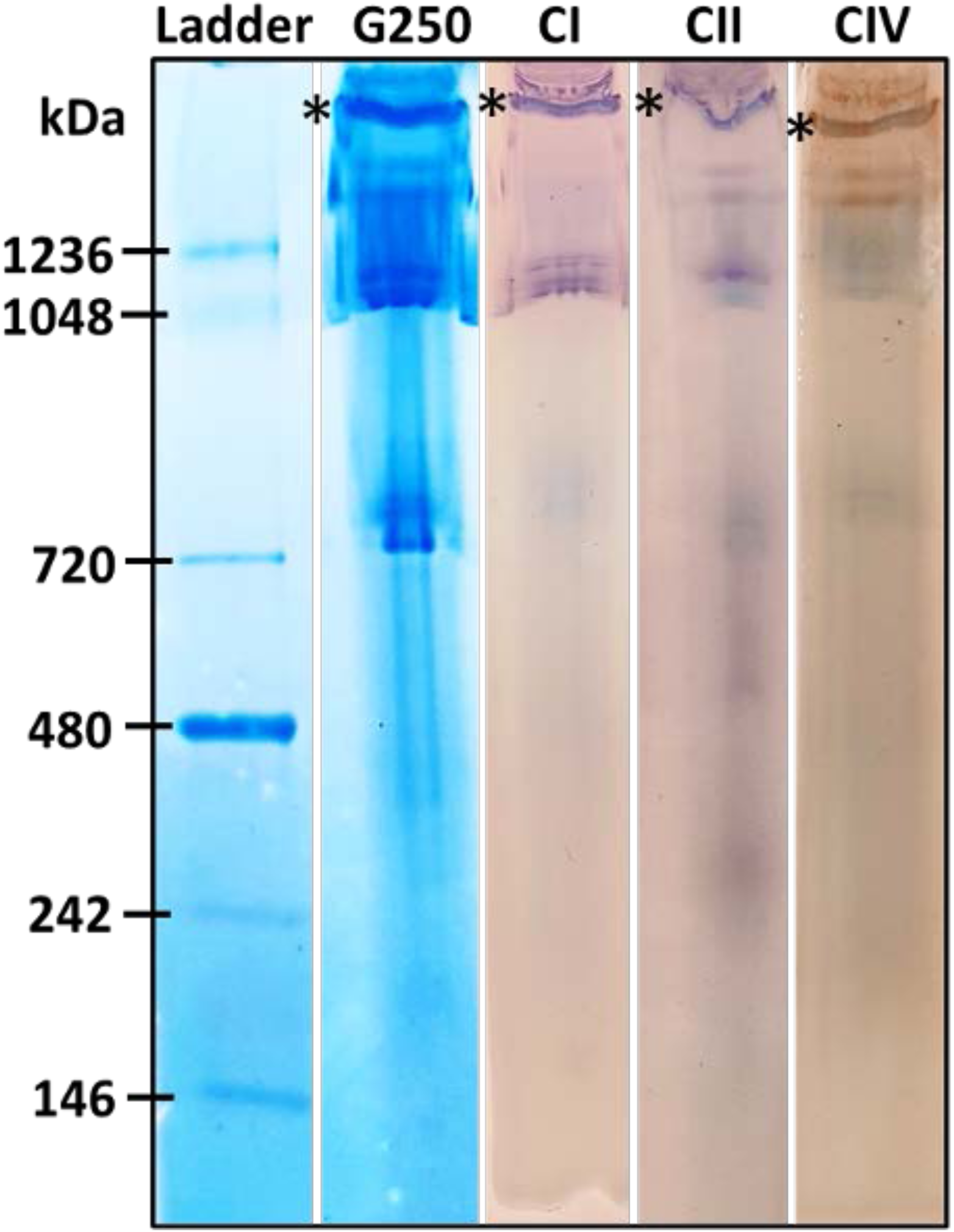
CN-PAGE gel showing active CI, CII and CIV in the final supercomplex sample. Coomassie stained CN-PAGE and in-gel activity assay to visualize the purified I-II-III_2_-IV_2_ supercomplex. CN-PAGE was performed to separate protein assemblies within the final sucrose cushion sample. The ladder and G250 lanes represent the Coomassie stained molecular weight marker and final supercomplex sample, respectively. CI, CII and CIV lanes highlight those protein bands with active CI, CII (purple) and CIV (brown), respectively. The band marked with an asterisk (*) was tentatively assigned as the intact supercomplex.

**Supplementary Information Figure 3.**
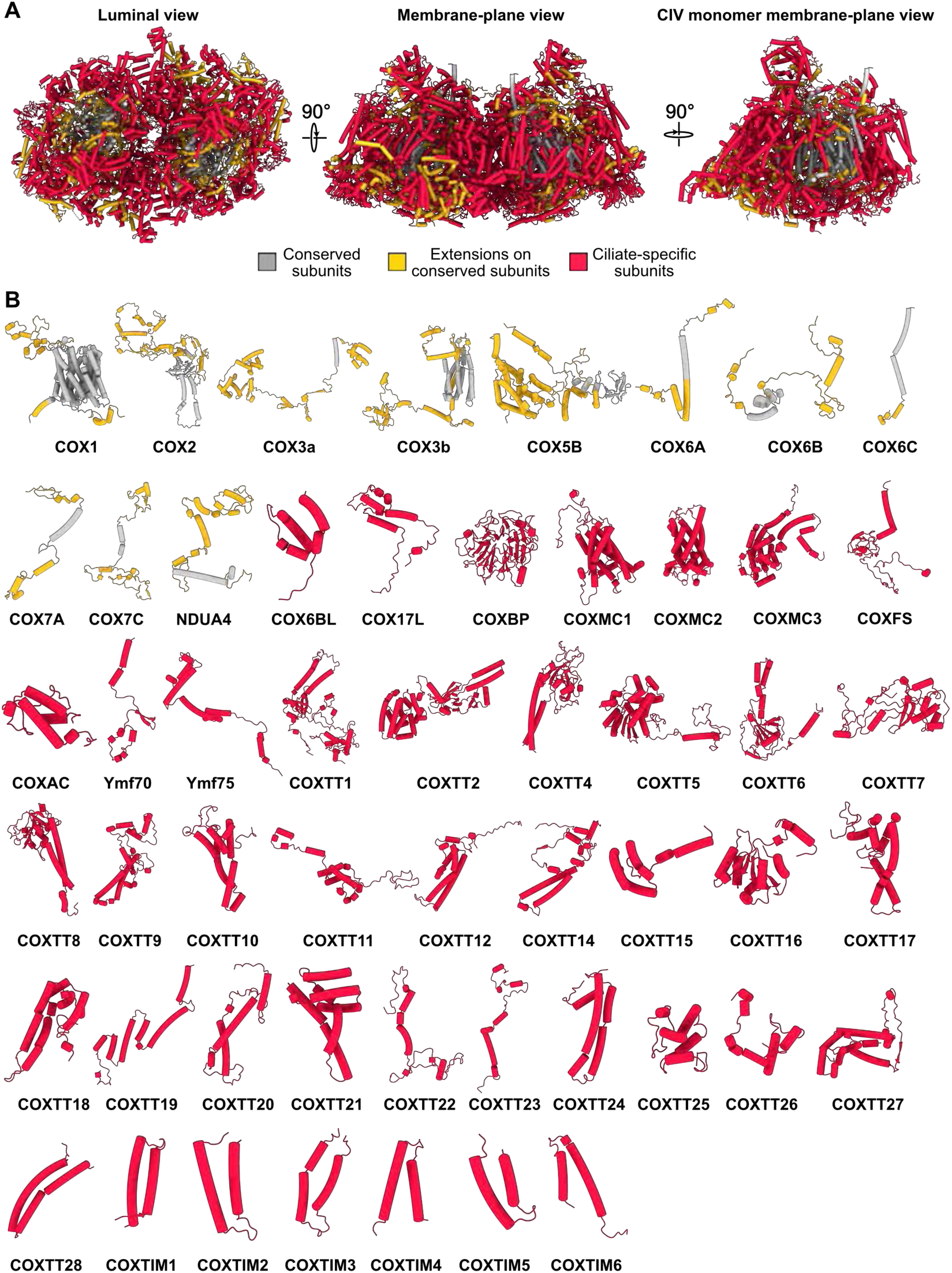
Conservation and divergence of ciliate CIV. **(A)** three different views of CIV_2_ colored by conserved subunits (grey), extensions to conserved subunits (gold) and ciliate-specific subunits (red). Conservation is assessed as structural homology to bovine (PDB 3X2Q) and human (PDB 5Z62) CIV structures. Right view shows a single CIV monomer assembly looking from the dimer interface. The ciliate CIV displays conserved core subunits that all contain significant extensions to the core folds. The conserved core subunits are also surrounded by ciliate-specific subunits, which make up most of the peripheral regions of the complex. **(B)** Individual CIV subunits shown as gallery with similar color coding as in A. Subunit naming is adopted from ref (*7*).

**Supplementary Information Figure 4.**
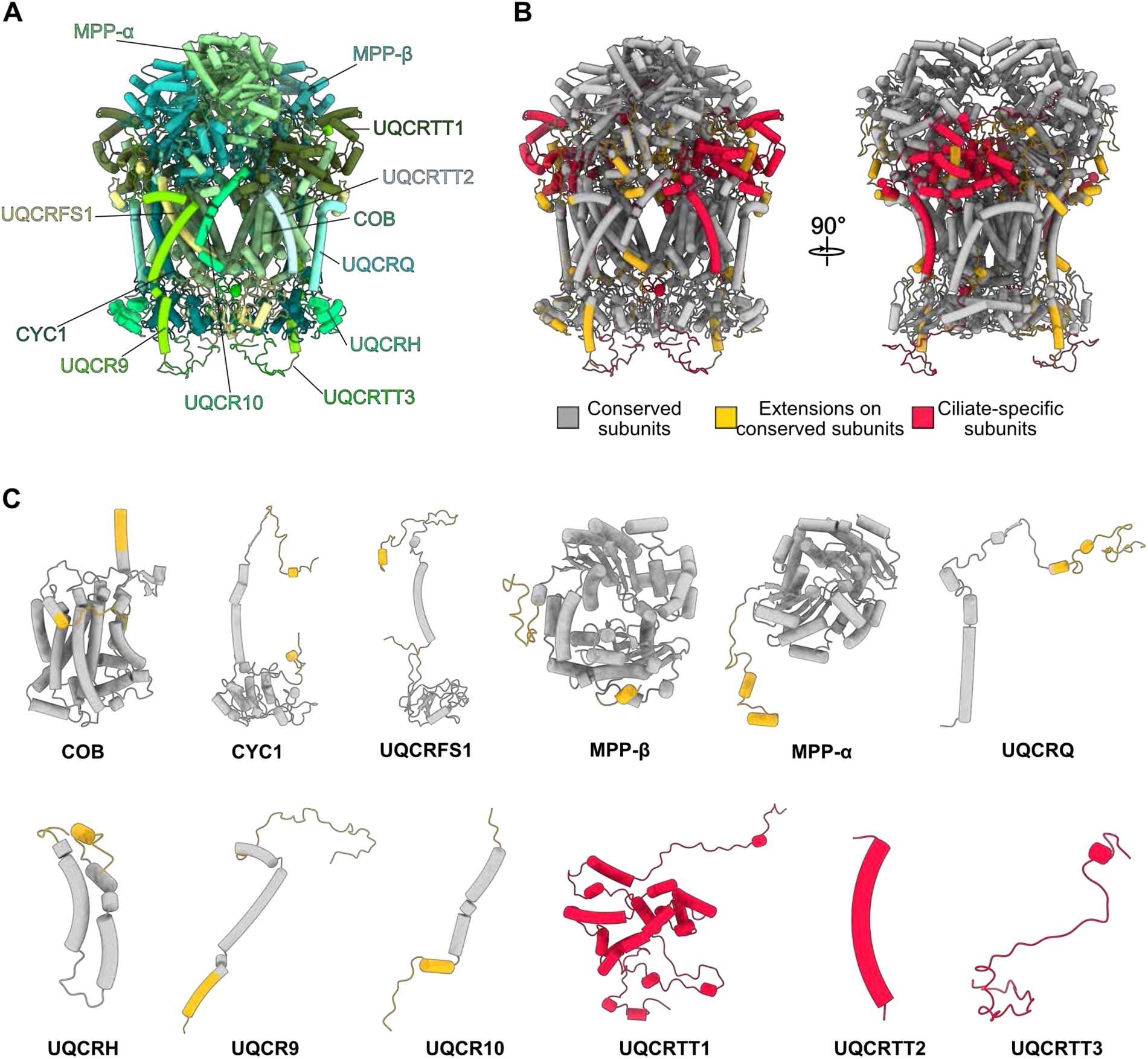
Conservation and divergence of ciliate CIII dimer. **(A)** CIII_2_ colored by subunits. **(B)** CIII_2_ colored by conserved subunits (grey), extensions to conserved subunits (gold) and ciliate-specific subunits (red). Conservation is assessed as structural homology to bovine (PDB 5J4Z) and murine (PDB 7O3H) CIII_2_ structures. The ciliate CIII_2_ shows a high degree of conservation in the core subunits. **(C)** Individual CIII subunits shown as gallery with similar color coding as in B.

**Supplementary Information Figure 5.**
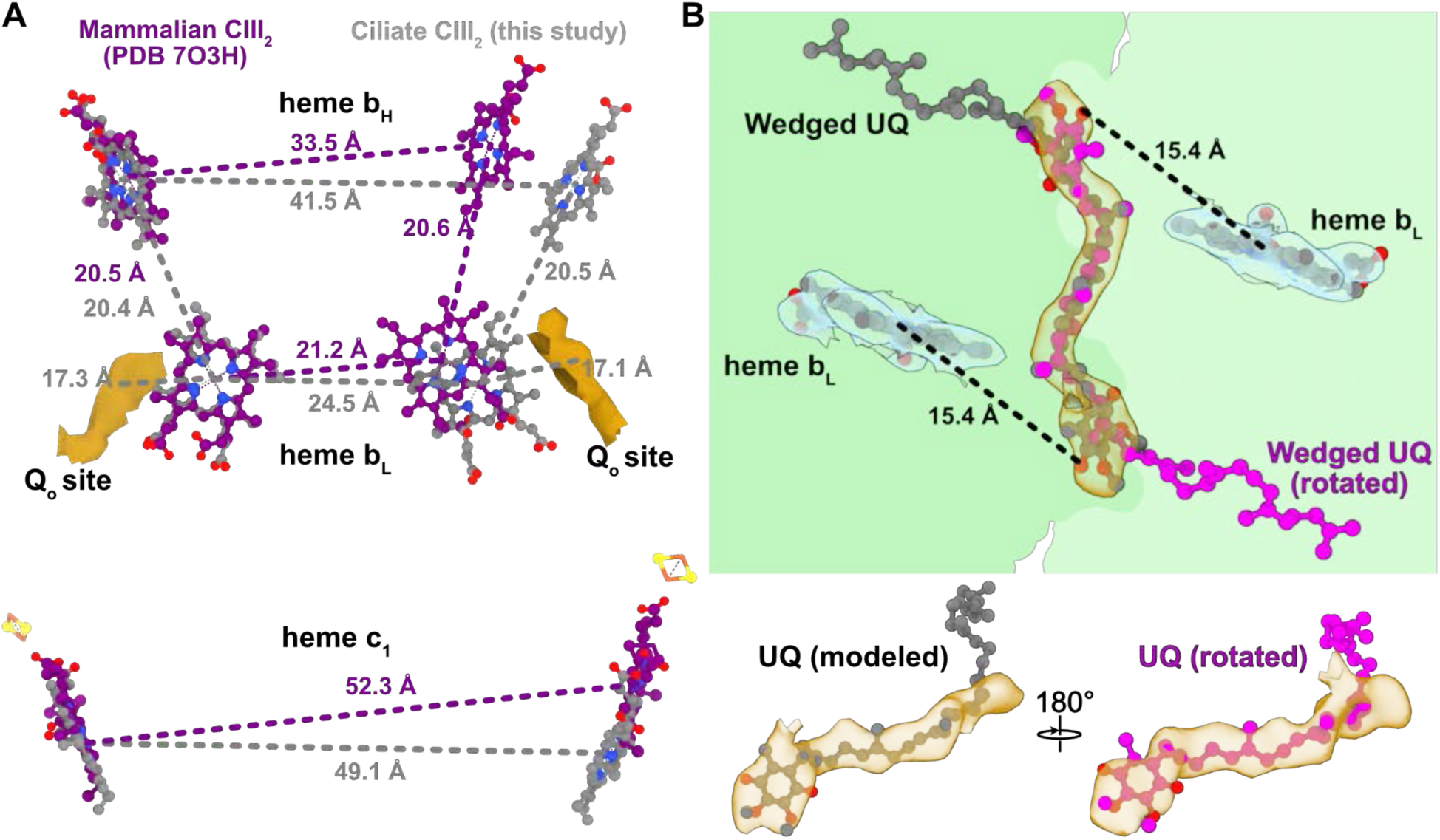
Ciliate CIII_2_ heme group distances and wedged ubiquinone. **(A)** Distance measures between ciliate (grey dashed lines) and mammalian (purple dashed lines) heme b_H_, heme b_L_ and heme c_1_ groups (measuring from Fe atoms) and the ciliate Q_o_ sites. Superposing mammalian CIII_2_ (PDB 7O3H) on one COB chain, shows that ciliate heme b_H_ and heme b_L_ (grey sticks) are displaced further away from the symmetry-related heme groups compared to mammalian, whereas the ciliate heme c_1_ groups are slightly closer. However, the distance between heme b_H_ and heme b_L_ within the same COB copy is essentially identical between ciliate and mammalian structures. Furthermore, the distance from the Q_o_ sites (middle of dark gold map density) to the heme b_L_ groups is comparable to the 16.8 Å distance between ubiquinol and the heme b_L_ group observed in the X-ray structure of bovine CIII_2_ (17). **(B)** Map density (transparent gold) of the CIII_2_ wedged ubiquinone (UQ) in COB (green) is almost rotationally symmetric around the dimer axis, showing planar density features for placement of UQ head groups equally close to the heme b_L_ molecules (transparent blue). We placed UQ in the orientation where the quinone fitted the map density best (grey) compared to the rotated orientation (magenta).

**Supplementary Video 1. Coarse-grained molecular dynamics simulation of the *T. thermophila* supercomplex.** First 800 ns of the MD simulation starting from an initially planar membrane reveals a deformation of the bilayer into a curved topology to accommodate the membrane protein complex.

**Supplementary Video 2. Annular lipid shell of the *T. thermophila* supercomplex.** Final frame of the coarse-grained MD-simulation with supercomplex and surrounding annular lipids shown, highlighting the curved shape of the lipid belt.

**Supplementary Information Table 1.** Data collection and model statistics

| Data collection | CI | CII | CIII<sub>2</sub> | CIV<sub>2</sub> | CI-II-III<sub>2</sub>-IV<sub>2</sub> |
| --- | --- | --- | --- | --- | --- |
| Microscope | Titan Krios |  |  |  |  |
| Voltage (kV) | 300 |  |  |  |  |
| Camera | K2 Summit |  |  |  |  |
| Magnification | 165,000 |  |  |  |  |
| Exposure (e<sup>-</sup>/Å<sup>2</sup>) | 30.9 |  |  |  |  |
| Pixel size (Å) | 0.83 |  |  |  |  |
| Defocus range (µm) | 0.6-2.6 |  |  |  |  |
| Movies collected | 26,063 |  |  |  |  |
| Frames / movie | 20 |  |  |  |  |
| Data processing |  |  |  |  |  |
| Initial particles | 1,664,103 |  |  |  |  |
| Final particles | 138,746 |  |  |  |  |
| Symmetry | C1 | C1 | C1 | C1 | C1 |
| Map resolution (Å) | 2.9 | 2.8 | 2.8 | 2.6 | 2.9 |
| - FSC threshold | 0.143 | 0.143 | 0.143 | 0.143 | 0.143 |
| B-factor sharpening | 46.8 | 49.1 | 52.2 | 45.4 | 49.9 |
| EMDB ID | EMD- | EMD- | EMD- | EMD- | EMD- |
| Model refinement statistics |  |  |  |  |  |
| CC (map/model) | 0.82 | 0.82 | 0.81 | 0.84 | 0.79 |
| Resolution (map/model) | 2.99 | 3.3 | 3.18 | 2.89 | 3.36 |
| - FSC threshold | 0.5 | 0.5 | 0.5 | 0.5 | 0.5 |
| Atoms (not H) | 234277 | 41399 | 94264 | 436066 | 806006 |
| Residues | 13712 | 2409 | 5438 | 23792 | 45351 |
| Lipids/CoQ | 59/0 | 11/1 | 32/3 | 214/8 | 316/12 |
| Model B-factor (Å<sup>2</sup>) |  |  |  |  |  |
| - Protein | 53.2 | 100.46 | 65.66 | 52.8 | 91.23 |
| - Ligands | 63.83 | 107.88 | 78.66 | 72.16 | 111.10 |
| Rotamer outliers (%) | 0.51% | 0.62% | 0.53% | 1.18% | 0.76 |
| Ramachandran (%) |  |  |  |  |  |
| - Outliers | 0.01% | 0.21% | 0.00% | 0.01% | 0.02% |
| - Allowed | 2.07% | 3.28% | 2.43% | 1.75% | 1.77 |
| - Favoured | 97.92% | 96.51% | 97.57% | 98.25% | 98.22 |
| Clash score | 4.12 | 3.91 | 5.68 | 4.22 | 4.63 |
| MolProbity score | 1.21 | 1.41 | 1.40 | 1.26 | 1.24 |
| RMSD |  |  |  |  |  |
| - Bonds (Å) | 0.002 | 0.004 | 0.002 | 0.002 | 0.002 |
| - Angles (°) | 0.407 | 0.459 | 0.447 | 0.503 | 0.562 |
| PDB ID |  |  |  |  |  |

**Supplementary Information Table 2.** List of proteins and comments

| Subunit name | Gene name | UniProt ID | Encoded | Residues in total | Comments |
| --- | --- | --- | --- | --- | --- |
| **Complex-I:** |  |  |  |  |  |
| ND1a | nad1_a | Q950Y3 | mt | 284 | Corresponds to NuoH, Nqo8 |
| ND1b | nad1_b | NP_149380 | mt | 59 | Mt-ND1 C-terminal extension, separate gene |
| ND2a | ymf65 | Q951A3 | mt | 360 | Corresponds to NuoN, Nqo14 |
| ND2b | nad2 | Q951B2 | mt | 178 | Mt-ND2 C-terminal extension, separate gene |
| Mt-ND3 | nad3 | Q950Z7 | mt | 121 | Corresponds to NuoA, Nqo7 |
| ND4 | nad4 | Q950X9 | mt | 505 | Corresponds to NuoM, Nqo13 |
| MT-ND4L | ymf58 | Q950Z5 | mt | 116 | Corresponds to NuoK, Nqo11 |
| ND5a | nad5 | Q950Z0 | mt | 750 | Corresponds to NuoL, Nqo12 |
| ND5b | ymf57 | Q951C2 | mt | 100 | Mt-ND5 C-terminal extension, separate gene |
| ND6 | ymf62 | Q950Y2 | mt | 255 | Corresponds to NuoJ, Nqo10 |
| NDUV1 | TTHERM_00193910 | Q23KE4 | nuclear | 474 | Corresponds to NuoF, Nqo1 |
| NDUV2 | TTHERM_00335630 | I7MEP0 | nuclear | 274 | Corresponds to NuoE, Nqo2 |
| NDUS1 | TTHERM_00194<br>260 | Q23KA9 | nuclear | 718 | Corresponds to NuoG,<br>Nqo3 |
| NDUS2 | nad7 | Q951B1 | mt | 442 | Corresponds to NuoD,<br>Nqo4 |
| NDUS3 | nad9_2 | Q950Z3 | mt | 198 | Corresponds to NuoC,<br>Nqo5 |
| NDUS4 | TTHERM_00526<br>930 | I7MK61 | nuclear | 185 | Corresponds to AQDQ<br>subunit |
| NDUS5 | TTHERM_00012<br>9647 | W7X4R4 | nuclear | 94 |  |
| NDUS6 | TTHERM_00497<br>570 | I7MK02 | nuclear | 132 | Zinc-finger protein,<br>contains a Zn<sup>2+</sup> ion |
| NDUS7 | nad10 | Q951B4 | mt | 162 | Corresponds to NuoB,<br>Nqo6 |
| NDUS8 | TTHERM_00294<br>640 | I7MDW5 | nuclear | 236 | Corresponds to NuoI,<br>Nqo9 |
| NDUA1 | TTHERM_00455<br>560 | I7MI60 | nuclear | 94 |  |
| NDUA2 | TTHERM_00659<br>070 | I7MA77 | nuclear | 103 |  |
| NDUA3 | TTHERM_00148<br>710 | I7M9B3 | nuclear | 135 |  |
| NDUA5 | TTHERM_01161<br>000 | Q23ND5 | nuclear | 206 |  |
| NDUA6 | TTHERM_00933<br>070 | I7M2Y3 | nuclear | 172 |  |
| NDUA7 | TTHERM_00399<br>360 | I7MIJ7 | nuclear | 282 |  |
| NDUA8 | TTHERM_00464<br>930 | I7MMF4 | nuclear | 238 |  |
| NDUA9 | TTHERM_00557<br>760 | I7MLH2 | nuclear | 362 | NAD-dependent<br>epimerase/dehydratase<br>family protein |
| NDUA11 | TTHERM_00945<br>210 | Q24F24 | nuclear | 213 |  |
| NDUA12 | TTHERM_00194<br>499 | A4VDQ6 | nuclear | 194 |  |
| NDUA13 | TTHERM_00149<br>260 | I7M2U4 | nuclear | 175 |  |
| NDUAB1<br>-α | TTHERM_01005<br>100 | Q22XT6 | nuclear | 138 | Different isoform to<br>NDUAB1-β |
| NDUAB1<br>-β | TTHERM_00470<br>710 | I7MD12 | nuclear | 133 | Acyl carrier protein |
| NDUB3 | TTHERM_00446<br>569 | A4VD20 | nuclear | 83 |  |
| NDUB4 | TTHERM_00310<br>880 | I7MG29 | nuclear | 126 |  |
| NDUB6 | TTHERM_00430<br>000 | Q231G0 | nuclear | 129 |  |
| NDUB7 | TTHERM_00402<br>070 | I7MIM0 | nuclear | 120 |  |
| NDUB8 | TTHERM_00481<br>330 | I7M855 | nuclear | 207 |  |
| NDUB9 | TTHERM_00985<br>010 | Q233X7 | nuclear | 189 |  |
| NDUB10 | TTHERM_00193<br>750 | Q23KG0 | nuclear | 188 |  |
| NDUB11 | TTHERM_00114<br>380 | Q22Z32 | nuclear | 214 |  |
| NDUB15 | TTHERM_00061<br>4679 | - | nuclear | 147 | Wrong gene annotation –<br>change translation ORF<br>to 5'-3' in frame 3 |
| NDUC2 | TTHERM_00160<br>690 | Q22W63 | nuclear | 102 |  |
| NDUCA1 | TTHERM_00136<br>440 | I7M8Q7 | nuclear | 346 | carbonic anhydrase-like<br>protein (isoform 1) |
| NDUCA2 | TTHERM_01005<br>010 | Q22XU5 | nuclear | 257 | carbonic anhydrase-like<br>protein (isoform 2) |
| NDUCA3 | TTHERM_00541<br>460 | I7M6S0 | nuclear | 233 | carbonic anhydrase-like<br>protein (isoform 3) |
| NDUX1 | TTHERM_00715<br>850 | I7LT42 | nuclear | 150 |  |
| NDUFX | TTHERM_00161<br>210 | Q22W11 | nuclear | 172 | 2Fe2S cluster-binding<br>protein |
| NDUTX | TTHERM_01205<br>250 | Q22AI5 | nuclear | 166 | Thioredoxin, TRX family |
| NDUJ1 | TTHERM_00938<br>750 | Q22DR7 | nuclear | 317 | DnaJ domain protein |
| NDUPH1 | TTHERM_00697<br>370 | Q24C39 | nuclear | 251 |  |
| NDUPH2 | TTHERM_00193<br>950 | Q23KE0 | nuclear | 189 |  |
| NDUTT1 | TTHERM_00925<br>340 | Q22E24 | nuclear | 516 | lipid A-disaccharide<br>synthase |
| NDUTT2 | TTHERM_01000<br>190 | Q24HK5 | nuclear | 333 | Acyl-CoA synthetase<br>(AMP-forming)/AMP-acid ligase II |
| NDUTT3 | TTHERM_00268<br>000 | I7LUQ4 | nuclear | 311 |  |
| NDUTT4 | TTHERM_00295<br>430 | I7MIE0 | nuclear | 212 |  |
| NDUTT5 | TTHERM_00649<br>080 | I7LT77 | nuclear | 205 |  |
| NDUTT6 | TTHERM_00334<br>340 | I7M1N8 | nuclear | 144 |  |
| NDUTT7 | TTHERM_00637<br>590 | Q22HE4 | nuclear | 143 |  |
| NDUTT8 | TTHERM_00006<br>120 | Q22SC4 | nuclear | 135 |  |
| NDUTT9 | TTHERM_00653<br>670 | Q23B10 | nuclear | 136 |  |
| NDUTT1<br>0 | TTHERM_00616<br>320 | I7MAF0 | nuclear | 127 |  |
| NDUTT1<br>1 | TTHERM_00185<br>570 | Q22T55 | nuclear | 113 |  |
| NDUTT1<br>2 | TTHERM_00835<br>330 | Q22E95 | nuclear | 93 |  |
| NDUTT1<br>3 | TTHERM_00992<br>800 | Q22DC2 | nuclear | 73 |  |
| NDUTT1<br>4 | TTHERM_00399<br>460 | I7MIK1 | nuclear | 71 |  |
| NDUTT1<br>5 | TTHERM_00063<br>7389 | - | nuclear | 237 | Wrong gene annotation –<br>change translation ORF<br>to 3'-5' in frame 1<br>Binds ADP-Mg<sup>2+</sup> |
| NDUTT1<br>6 | TTHERM_00124<br>3407 | W7XFJ5 | nuclear | 119 | Is required for I-IV<sub>2</sub><br>interface stability.<br>Stabilises ND5a Nt<br>extension. |
| NDUTT1<br>7 | TTHERM_00053<br>2499 | - | nuclear | 125 | Wrong gene annotation –<br>change translation ORF<br>to 3'-5' in frame 2 |
| **Complex-**<br>**II:** |  |  |  |  |  |
| SDHA | TTHERM_00047<br>080 | Q23DI3 | nuclear | 636 |  |
| SDHB | TTHERM_00241<br>700 | I7M403 | nuclear | 312 |  |
| SDHC | TTHERM_00387<br>120 | Q23RH8 | nuclear | 60 |  |
| SDHD | AF396436.1 (mt-<br>genome<br>sequence) | - | mt | 43 | Gene lies between ymf66<br>and ymf76 on mtDNA.<br>Translation 3'-5' in frame<br>1 |
| SDHTT1 | TTHERM_00571<br>650 | Q24I09 | nuclear | 322 | contains heme c |
| SDHTT2 | TTHERM_00532<br>090 | Q248F8 | nuclear | 296 |  |
| SDHTT3 | TTHERM_00283<br>850 | I7MEX7 | nuclear | 198 |  |
| SDHTT4 | TTHERM_00658<br>950 | I7LX66 | nuclear | 195 | Diphthamide synthesis<br>protein |
| SDHTT5 | TTHERM_00601<br>860 | Q22YL0 | nuclear | 114 |  |
| SDHTT6 | TTHERM_00626<br>980 | Q23S01 | nuclear | 103 |  |
| SDHTT7 | TTHERM_00713<br>350 | Q24CW6 | nuclear | 93 |  |
| SDHTT8 | TTHERM_00028<br>7919 | W7XBF5 | nuclear | 89 |  |
| SDHTT9 | TTHERM_00637<br>670 | Q22HD6 | nuclear | 76 |  |
| SDHTT10 | TTHERM_00103<br>4353 | W7XF00 | nuclear | 62 |  |
| SDHTT11 | DY684362.1<br>(EST, mRNA<br>sequence) | - | nuclear | 46 | Gene translation should<br>be 3'-5' in frame 1 |
| **Complex-**<br>**III:** |  |  |  |  |  |
| MPP-β | TTHERM_00502<br>380 | I7MGU2 | nuclear | 513 |  |
| MPP-α | TTHERM_00836<br>690 | I7MJ25 | nuclear | 482 |  |
| COB | cob | Q950Z1 | mt | 426 | contains heme bL and bH |
| CYC1 | TTHERM_00918<br>500 | Q24IM5 | nuclear | 319 | Contains heme c |
| UQCRFS<br>1 | TTHERM_00295<br>080 | I7MIC7 | nuclear | 269 | Contains 2Fe2S cluster |
| UQCRH | TTHERM_00194<br>690 | Q23K66 | nuclear | 86 | cytochrome bc1 hinge<br>proteins |
| UQCRQ | TTHERM_00765<br>330 | I7M484 | nuclear | 130 |  |
| UQCR9 | TTHERM_00456<br>790 | I7MM45 | nuclear | 119 |  |
| UQCR10 | TTHERM_00218<br>930 | I7MFL6 | nuclear | 62 |  |
| UQCRTT<br>1 | TTHERM_00382<br>330 | Q23F81 | nuclear | 328 |  |
| UQCRTT<br>2 | EV837551.1<br>(EST, mRNA<br>sequence) | - | nuclear | 41 | Gene translation should<br>be 5'-3' in frame 3 |
| UQCRTT<br>3 | - | - | - | 66 | Unidentified subunit |
| **Complex-**<br>**IV:** |  |  |  |  |  |
| COX1 | cox1 | Q950Y4 | mt | 688 |  |
| COX2 | cox2 | Q950Y9 | mt | 604 |  |
| COX3a | ymf67 | Q950Y7 | mt | 453 | structurally replaces H1 of COX3 |
| COX3b | ymf68 | Q950Y6 | mt | 594 |  |
| COX5B | TTHERM_00378<br>620 | Q23FF5 | nuclear | 637 |  |
| COX6A | TTHERM_00072<br>3218 | W7XCY<br>5 | nuclear | 130 |  |
| COX6B | TTHERM_00568<br>030 | Q24I72 | nuclear | 230 |  |
| COX6C | TTHERM_00046<br>170 | Q23DS4 | nuclear | 103 |  |
| COX7A | TTHERM_00151<br>250 | I7MGF9 | nuclear | 133 |  |
| COX7C | TTHERM_00047<br>0561 | W7X287 | nuclear | 236 |  |
| COX6BL | TTHERM_00218<br>340 | I7LVX0 | nuclear | 88 |  |
| COX17L | TTHERM_01043<br>280 | Q22CI1 | nuclear | 990 | CTF/NF-I domain-containing protein |
| NDUA4 | TTHERM_00100<br>1528 | W7WZP<br>1 | nuclear | 220 | (Named COXTT13 in Zhou et al.) |
| COXMC1 | TTHERM_00127<br>269 | A4VDV3 | nuclear | 346 | Oxoglutarate/malate translocator protein |
| COXMC2 | TTHERM_00621<br>600 | Q23M99 | nuclear | 318 | 2-oxoglutarate/malate carrier protein |
| COXMC3 | TTHERM_00112<br>650 | Q22ZA6 | nuclear | 330 |  |
| COXBP | TTHERM_00016<br>360 | Q22RF2 | nuclear | 685 | BBC53 chromosome<br>condensation regulator<br>RCC1 |
| COXTIM<br>1 | TTHERM_00052<br>8460 | W7X3D6 | nuclear | 72 | Tim10/DDP family zinc<br>finger proteins |
| COXTIM<br>2 | TTHERM_00070<br>3379 | W7XDM<br>6 | nuclear | 72 | part of TIM9.10 hexamer |
| COXTIM<br>3 | TTHERM_00433<br>490 | Q231A8 | nuclear | 93 | part of TIM9.10 hexamer |
| COXTIM<br>4 | TTHERM_01289<br>060 | Q22A35 | nuclear | 68 | part of TIM9.10 hexamer |
| COXTIM<br>5 | TTHERM_00028<br>510 | Q22N23 | nuclear | 81 | part of TIM9.10 hexamer |
| COXTIM<br>6 | TTHERM_00805<br>850 | Q233U0 | nuclear | 72 | part of TIM9.10 hexamer |
| COXFS | TTHERM_00133<br>510 | I7M8P0 | nuclear | 188 | Fe-binding zinc finger<br>CDGSH type protein |
| COXAC | TTHERM_00695<br>750 | Q24C97 | nuclear | 127 |  |
| Ymf70 | ymf70 | Q950Y0 | mt | 89 |  |
| Ymf75 | ymf75 | Q951A7 | mt | 190 |  |
| COXTT1 | TTHERM_00361<br>490 | Q22PJ5 | nuclear | 490 |  |
| COXTT2 | TTHERM_00721<br>790 | Q22FX8 | nuclear | 473 | Protein phosphatase 2C |
| COXTT4 | TTHERM_00338<br>280 | I7M1Q4 | nuclear | 402 | SURF1 family protein |
| COXTT5 | TTHERM_00046<br>440 | Q23DP7 | nuclear | 385 | TraB family protein |
| COXTT6 | TTHERM_00047<br>230 | Q23DG8 | nuclear | 348 |  |
| COXTT7 | TTHERM_00675<br>650 | Q23DZ5 | nuclear | 318 |  |
| COXTT8 | TTHERM_00070<br>850 | I7LTZ4 | nuclear | 318 | SURF1 family protein |
| COXTT9 | TTHERM_00525<br>160 | I7LY65 | nuclear | 252 |  |
| COXTT10 | TTHERM_00420<br>130 | I7MD70 | nuclear | 234 |  |
| COXTT11 | TTHERM_00093<br>9159 | W7X4J9 | nuclear | 231 |  |
| COXTT12 | TTHERM_00455<br>090 | I7M3P9 | nuclear | 215 |  |
| COXTT14 | TTHERM_00530<br>650 | I7LZX8 | nuclear | 210 |  |
| COXTT15 | TTHERM_00641<br>250 | Q23F08 | nuclear | 193 |  |
| COXTT16 | TTHERM_00218<br>840 | I7M8Y9 | nuclear | 175 | AAA protein fold |
| COXTT17 | TTHERM_00049<br>040 | Q23D87 | nuclear | 173 |  |
| COXTT18 | TTHERM_00218<br>570 | I7MKT6 | nuclear | 173 |  |
| COXTT19 | TTHERM_00433<br>830 | Q230X6 | nuclear | 170 |  |
| COXTT20 | TTHERM_00794<br>470 | Q23VY4 | nuclear | 158 |  |
| COXTT21 | TTHERM_00938<br>940 | Q22DP8 | nuclear | 154 |  |
| COXTT22 | TTHERM_00691<br>100 | I7MFV5 | nuclear | 149 |  |
| COXTT23 | TTHERM_00666<br>370 | Q23TE5 | nuclear | 124 |  |
| COXTT24 | TTHERM_00161<br>000 | Q22W32 | nuclear | 122 |  |
| COXTT25 | TTHERM_00151<br>580 | I7M9E7 | nuclear | 105 |  |
| COXTT26 | TTHERM_00485<br>790 | I7LTF1 | nuclear | 90 |  |
| COXTT27 | TTHERM_00030<br>2101 | W7XDH<br>2 | nuclear | 212 | (Including subunit<br>COXTT3 in Zhou et al.) |
| COXTT28 | TTHERM_00093<br>3397 | W7X912 | nuclear | 171 |  |

